# Derivation of trophoblast stem cells unveils unrestrained potential of mouse ESCs and epiblast

**DOI:** 10.1101/2023.04.19.537518

**Authors:** Debabrata Jana, Purnima Sailasree, Priya Singh, Mansi Srivastava, Vijay V Vishnu, Hanuman T Kale, Jyothi Lakshmi, Gunda Srinivas, Divya Tej Sowpati, P Chandra Shekar

## Abstract

mESCs and epiblast are considered to follow strict lineage adherence and lack the potential to contribute to trophoectoderm. Here, we report the derivation of trophoblast stem cells (ESTS) from the mESCs. The single-cell transcriptome and molecular characterization of ESTS show similarity with TSCs. They efficiently integrate into the TE compartment of the blastocyst and contribute to the placenta during development. We discovered GSK3β as a critical regulator of the TE fate of ESCs. It plays a vital stage-specific role during ESTS derivation. We further show β-CATENIN and an intron-I regulatory element of *Cdx2* are essential for the TE fate of ESCs. We further show that the mouse epiblast can readily differentiate into TE lineage. In contrast to the paradigm of the restricted potential of pluripotent ESCs and epiblast, our data shows that murine ESCs and epiblast have the unrestrained developmental potential for extraembryonic lineages.

## INTRODUCTION

The mammalian zygote undergoes cleavage followed by the segregation of an outer trophoectoderm (TE) layer and an inner cell mass (ICM) to form the blastocyst. The TE and its cell culture equivalent - trophoblast stem cells (TSCs) can contribute to the extraembryonic tissue -placenta. The ICM further delineates into the epiblast and the primitive endoderm(Cockburn and Rossant, 2010; Gardner and Rossant, 1979). The ESCs derived from the epiblast have restricted potential to contribute to the embryo and primitive endoderm derivatives(Chazaud et al., 2006; Gardner and Rossant, 1979; Plusa et al., 2008). They lack the potential to contribute to the TE derivatives(Gardner, 1983; Nichols and Gardner, 1984). The textbook model of strict adherence to the restricted lineage potential of the mammalian epiblast and ESCs is being challenged by recent reports of the derivation of TSCs from the human pluripotent stem cells (PSCs)(Dong et al., 2020; Guo et al., 2021; Jang et al., 2022; Wei et al., 2021). However, the ability of such cells to contribute placenta *in vivo* is unverified. Further, the lack of TSCs derived from PSCs of any other mammals continues to support the adherence to the restricted lineage potential of mammalian PSCs. The TSCs induced from the mouse pluripotent stem cells by genetic manipulations involving cellular reprogramming fail to attain the transcriptome and chromatin architecture of the TSC derived from a blastocyst(Cambuli et al., 2014; Kuckenberg et al., 2010; Ng et al., 2008; Nishioka et al., 2009; Niwa et al., 2000; Niwa et al., 2005; Ralston et al., 2010). Despite decades of research efforts, the derivation of TSCs from murine ESCs or epiblast has remained elusive leading to the conclusion of a lack the TE potential(Guo et al., 2021) in murine ESCs and epiblast.

Contrary to the current understanding, we report that the mouse ESCs and the mouse epiblast have the potential to contribute to TE lineage by derivation of TSCs. TSCs can be derived by the priming of ESCs to TE lineage and conversion to TSCs under defined culture conditions. The transcriptome and developmental potential of ESTS are similar to the TSCs. We identified GSKβ activity as the gatekeeper of the TE potential of ESCs. β-CATENIN and *Cdx2*-intron-I regulatory elements are essential for the TE potential of ESCs. We further show that mouse epiblast has the potential readily differentiate into TE lineage cells. We show that mouse ESC and epiblast have full developmental potential to differentiate into all extraembryonic lineages.

## RESULTS

### Derivation and characterisation of trophoblast stem cells (ESTS) from ESCs

While studying the regulation of core pluripotency factors by GSK3β inhibitor - CHIR99201 (CHIR) and MEK inhibitor PD325901 (PD) in ESC(Ying et al., 2008), we observed cobblestone-shaped cells differentiating around the ESCs in presence of CHIR in serum and LIF (SLCHIR). They morphologically resembled the TE cells differentiated from ZHBTc4 ES cells by doxycycline treatment (ZHBTc4+Dox)(Niwa et al., 2000) (Figure 1A) and expressed TE master regulator *Cdx2*. Other TE factors were induced at some time points of SLCHIR treatment, *Cdx2* transcripts were strongly induced with time in SLCHIR than other TE transcripts (Figure S1A). The expression of *Cdx2* is dependent on the concentration of CHIR, which is reduced by PD in a dose-dependent manner (Figure S1B). The *Cdx2* expression is much higher in SLCHIR than in SL2i. CDX2 protein was expressed in SLCHIR, multiple folds higher than SL2i (PD+CHIR) and ZHBTc4+Dox cells consistent with the transcript levels. GATA3 was detectable in SLCHIR albeit lower than ZHBTc4+Dox cells (Figure 1C, S1C). Immunofluorescence showed only a subpopulation of ESCs expressed CDX2 in SLCHIR and SL2i (Figure 1D). A subpopulation of these cells co-expressed OCT4 and CDX2. Such cells were more frequent in SL2i than in SLCHIR. CDX2 expression was observed in the cells in the periphery of the ESC colony (Figure S1D), where cells mostly primed for differentiation reside. We generated and utilized TCMC-OGFP cell line to understand the origin of these subpopulations. The *Cdx2* expression in TCMC-OGFP is reported by mCherry(Jana et al., 2019) and *Oct4* expression by GFP (Figure S1E). FACS analysis of the SLCHIR treatment time course showed that the OCT4-expressing cells gave rise to *Oct4*-*Cdx2* double-positive cells. *Cdx2*-mCherry positive cells appear 16hrs onwards (Figure S1F). Most double-positive cells differentiated and lost OCT4 expression when cultured in TS media(Tanaka, 2006). A small proportion of *Cdx2*-expressing cells continued to express *Cdx2* in TS media, which was lost in subsequent passage suggesting that CHIR induced *Cdx2* expression was transient and could not be sustained by self-activation (Figure S1G, H).

**Fig. 1.**
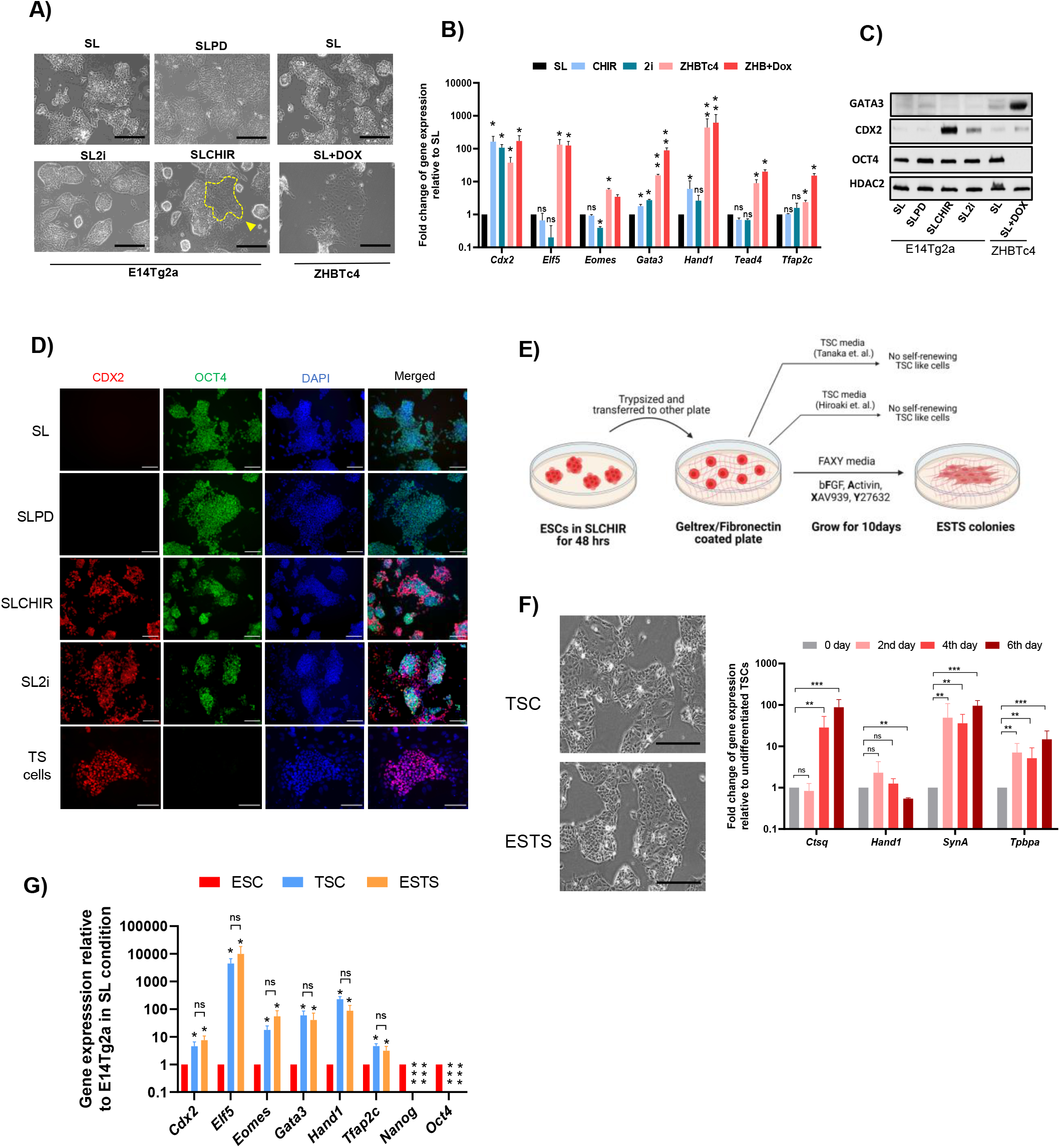
Derivation and characterization of ESTS from mouse ESC. A) Phase contrast images of E14Tg2a cultured in SL, SLPD, SLCHIR, SL2i, and ZHBTc4 cells cultured in the presence and absence of doxycycline (scale bar=100µm). TE-like cells in SLCHIR are marked with the yellow arrow. B) Quantitative expression analysis of TE lineage genes in indicated culture conditions, error bar represents SD of the mean of biological replicates (n=3). ‘*’ indicates a p-value <0.05, ‘**’ indicates a p-value <0.01, ‘ns’ indicates a p-value > 0.05. C). Immunoblot analysis of GATA3, CDX2, and OCT4 in indicated culture conditions of E14Tg2a and ZHBTc4 cells. D) Immunofluorescence analysis of CDX2 and OCT4 in E14Tg2a and TSCs derived from the blastocyst. Scale bar = 50µm. E) The experimental scheme depicting the derivation of ESTS from mouse ESCs. F) (Left) Phase contrast images of TSCs derived from the blastocyst and ES-TSCs derived from ESCs. Scale bar = 100µm. (Right) Relative mRNA abundance of TE differentiation genes analyzed by q-RTPCR in 0, 2^nd^, 4^th^, and 6^th^ day of differentiation. G) Relative mRNA abundance of TE and pluripotency genes analyzed by q-RTPCR. The error bar represents the SD of the mean of biological replicates (n=3). ‘*’ indicates a p-value <0.05, ‘**’ indicates a p-value <0.01, ‘***’ indicates a p-value <0.001, ‘ns’ indicates a p-value > 0.05.

*Cdx2* overexpression in ES cells represses *Oct4* to induce differentiation to TE and enable derivation of TSCs(Niwa et al., 2005). Transient *Cdx2* induction prompted us to derive TSC from SLCHIR-treated ESC. Our attempts to derive TSCs in TS media(Tanaka, 2006) and defined TSC media for human TSCs(Okae et al., 2018) from ESC treated with or no CHIR failed (Figure 1E, S2A). Although some cells with morphological resembles to TSCs were observed in both TS medias after SLCHIR treatment, they failed to self-renew in subsequent passages. Surprisingly, we could derive and maintain TSC-like cells in FAXY media (bFGF, Activin-A, XAV-939, and Y-27632)(Ohinata and Tsukiyama, 2014) from CHIR-treated ESC (Figure 1E, Figure S2A, B). ESC cultured in SL give rise to TSC-like cells in FAXY at very low frequency, suggesting that priming ESC with CHIR is essential to enhance the efficiency of TSC-like cells derivation (Figure S2A). These results suggest that CHIR partially primes a subpopulation of ESC towards TE lineage by inducing *Cdx2.* We refer to this state of ESC as TSC-primed-ESC. The cells might require further cues to make complete transition to TSCs which are lacking in TS media and human TSC media. However, in FAXY they make complete transition to TE lineage and enable derivation of TSC-like cells from ESC. The TSC-like cells derived from SLCHIR treated ESC were named as ESTS.

We supplemented FAXY with CHIR to increase ESTS derivation, surprisingly ESTS could not be derived in FAXY+CHIR (Figure S2C). CHIR promotes β-Catenin signalling while XAV-939(Huang et al., 2009) represses it, suggesting, β-Catenin signalling is essential for the priming stage but inhibitory for the next stage of transition to ESTS. This was further supported by reduced CDX2 and GATA3 expression when a lower concentration of XAV (5nM) was used instead of 10 nM in FAXY (Figure S2D). This data suggest that β-CATENIN has a stage-specific function during the derivation of ESTS. Further, the ESTS established in FAXY could be maintained and cultured for multiple passages in TS media. This suggests that although TS media may support self-renewal of ESTS, TS media may lack the ability to support the initial transition of the TSC-primed-ESC to TE lineage. The ESTS resembled TSCs derived from blastocysts (Figure 1F) and expressed CDX2 and GATA3 (Figure S2D, E). The ESTS could be passaged continuously and can differentiate to TE lineage (Figure 1F). We compared the expression of major TSC markers in ESTS. The expression of *Cdx2, Gata3, Elf5, Tfac2c, Eomes,* and *Hand1* in ESTS was comparable to TSCs (Figure 1G).

Together our data show that murine ESCs have the potential to contribute to TE lineage. GSK3β activity functions as a gatekeeper of TE lineage and its inhibition primes the ESCs to TE lineage and enable derivation of ESTS in FAXY media.

### Transcriptome analysis and developmental potential of ESTS

We carried out whole transcriptome analysis of the ESTSs, TSCs and ESCs. In a principal component analysis of the transcriptomes, the ESTSs clustered closer to TSCs suggesting they share similar transcriptome (Figure S3A). Expression of TE markers in ESTS were comparable to TSCs, and the pluripotency transcripts were absent in ESTS (Figure S3B-C). TSC genes like *Elf5, Tfap2c* and *Tead4* known to maintain self-renewal *were* relatively higher in ESTSs than in TSCs, while TSC differentiation gene like *Plet1*, *Krt7*, and *Hand1*(Marchand et al., 2011; Murray et al., 2016) were enriched in TSCs suggesting the self-renewal ability of ESTSs could be similar if not better than the TSCs (Figure S3B).

To ascertain the cellular identities, we performed the scRNA-Seq analysis of ESCs, TSCs, and ESTSs. The UMAP revealed an overlap of most ESTS and TSCs in clusters 0,1 and 3 in similar proportions. Cluster 2 & 4 represent mostly TSC and ESTS respectively but cluster closely. Together the data suggests that the ESTS are composed of cell types similar to TSCs (Figure 2A-B). Most ESCs are segregated into a distinct cluster 5. The TSCs are characterized by heterogenous culture similar to ESCs. TSC culture is composed of coexistence of multiple developmental transition stages. Most TSC gene like *Sox21*, *Elf5*, *BMP4*, *Sbsn*, *Bmp8b*, *Nat8l* and *Wnt6*(Frias-Aldeguer et al., 2019; Jaber et al., 2022; Li et al., 2013; Ralston et al., 2010; Zita et al., 2015) are enriched in cluster 0 (Figure 2C, S3C). The *Elf5* is repressed by DNA methylation in ESCs, the expression of *Elf5* and other polar TE genes like *Ppp2r2c*, *Eomes*, *Cdx2*, *Ddah1*, *Duox2*, *Cpne3* and *Rhox5*(Frias-Aldeguer et al., 2019) are enriched more in clusters 0, 2 and 4 (Figure S3D), suggesting these cluster may represent polar TE subpopulations (Figure S2D). Mural TE markers are distributed across except *Flt1*, *Slc5a5*, and *Slc2a3,* which are enriched partially in other clusters. Together our data suggest that the ESTS have heterogeneity similar to the TSCs and they are composed of large sub-populations of cells with expression of polar TE markers.

**Fig. 2.**
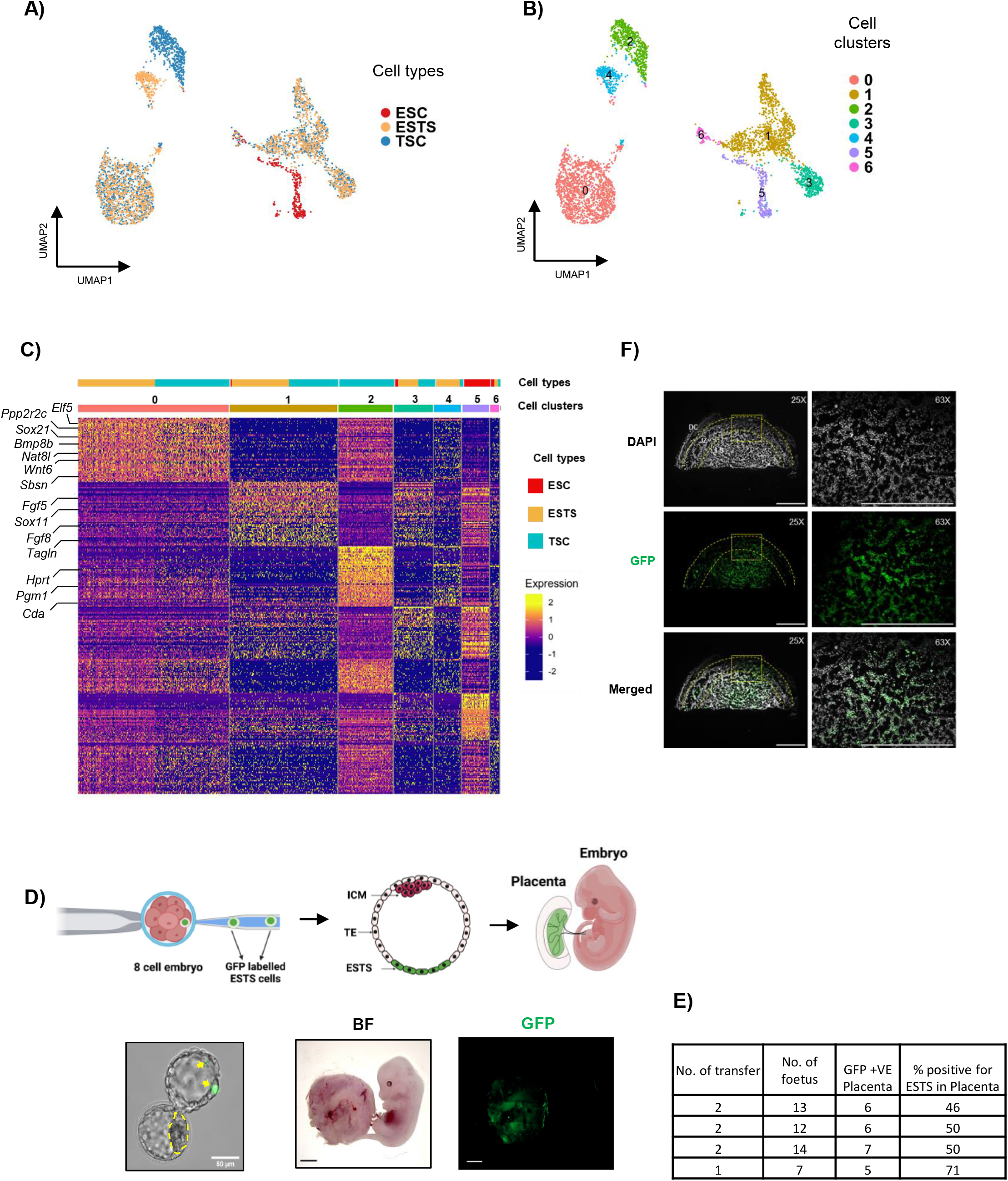
Single-cell transcriptomics of ESTS and its characterization. A) UMAP projection of ESC, TSC, and ESTS from sc-RNA Seq analysis. B) Cluster analysis of the UMAP projection of ESC, TSC, and ESTS from sc-RNA Seq analysis. C) Heatmap of the top 50 genes differentially expressed in different clusters of ESC, TSC, and ESTS from sc-RNA seq. D) (Top) Schematic showing injection of GFP labelled ESTS (ESTS-GFP) into 8-cell stage embryo and contribution of ESTS in blastocyst and placenta. (Left bottom) Representative image showing the contribution of GFP labelled ESTS in the TE layer of the blastocyst (marked with yellow arrow). (Right bottom) Representative image showing the contribution of GFP labelled ESTS descendant cells to the placenta at E12.5 (Fluorescence channel was acquired in tile scan). Scale bar =10mm. E) Summary of the contribution of GFP labelled ESTS to the placenta at E12.5. F) Immunofluorescence assay for detection of GFP-positive ESTS in the placental section. DC, JZ and LB depict the decidual region, junctional zone, and labyrinth region of the placenta respectively. Scale bar = 2000µm.

To exclude the possibility of cross-contamination of TSCs in ESTS culture, we derived ESTS from an ES cell line carrying Ef1a-H2B GFP transgene. ESTS-GFP cells (2 cells) were injected into morula stage embryos to assess their developmental potential. GFP-expressing cells were observed exclusively in the TE layer of the blastocysts. The GFP-expressing cells also expressed TE marker – GATA3 (Figure 2D, S3F). We transferred the blastocysts into the uterus of pseudo-pregnant mice and analysed the contribution of ESTS-GFP cells in E12.5 embryos. The ESTS-GFP cells contributed to the placenta, GFP expressing cells were not detected in the embryos (n=46) (Fig.2D-E, S3G). ESTS contributed to the placenta in nearly 52% of the chimeric embryos in agreement with contribution reported for TSCs earlier(Cambuli et al., 2014). ESTS-GFP contributed to the labyrinth as well as the junctional zone of the chimeric placenta (Figure 2F, S3H), suggesting ESTS contribute exclusively to TE lineage in developing chimeric embryos. Our data demonstrates that the ESTSs are heterogenous culture with subpopulations sharing transcriptome similar to TSCs. They can integrate into embryos to form chimeric embryos, participate in devolvement and efficiently contribute to the TE lineage derivatives.

### β-CATENIN and Intron-I regulatory region of *Cdx2* are essential for priming of ESC to TSCs

OCT4 and NANOG repress *Cdx2* to inhibit TE lineage(Chen et al., 2009; Niwa et al., 2000). We asked whether *Nanog* or *Oct4* expression levels affect CHIR-induced expression of *Cdx2* and priming of ESCs to TE during ESTS derivation. We utilised *Nanog*:GFP (TNGA)(Chambers et al., 2007) and *Oct4*:GFP (Oct4GiP)(Kale et al., 2022) ESC lines to sort the lowest 10% and highest 10% *Nanog* or *Oct4* expressing cells. The *Cdx2* transcript was analysed after 16hrs of CHIR treatment. *Cdx2* was induced significantly in low-*Nanog* relative to high*-Nanog* population but remained unchanged in *Oct4*-low and high cells, suggesting CHIR induces *Cdx2* in low-*Nanog* cells (Figure 3A). Further, we utilised genetically engineered ESC with different levels of NANOG and OCT4 to analyse the induction of *Cdx2* (Figure 3B). CDX2 was detectable in the presence of CHIR in all the cell lines. Constitutive expression of *Nanog* transgene in the ENOE cell line significantly repressed CDX2. CDX2 was induced multiple folds in *Nanog* null cell line and was barely detectable when NANOG expression was restored from a transgene in TBCR cell line (Figure 3C, S4A) suggesting that NANOG prepresses CHIR mediated induction of CDX2 in a dosage dependent manner. CDX2 was induced in *Oct4*+/- cell in CHIR, surprisingly, it was further increased in cell line with constitutive over-expression of *Oct4*, suggesting that the basal expression levels of OCT4 are essential and sufficient to repress *Cdx2* in ESC (Figure 2C, S4B). However, OCT4 does not show dosage-dependent repression of *Cdx2,* unlike NANOG. To understand how OCT4 and NANOG repress CHIR-induced *Cdx2* expression, we utilised ZHBTc4 cells where *Oct4* could be shut down by Doxycycline to induce differentiation to TE lineage(Niwa et al., 2000). We constitutively over-expressed Flag-Bio-*Nanog* from the transgene in ZHBTc4 to generate ZHBNOE (Figure S4C). Doxy treatment induced expression of CDX2 and GATA3 in ZBHTc4 and ZHBNOE (Figure 3D). Overexpression of NANOG in ZHBNOE failed to repress CDX2 in absence of OCT4 resulting in NANOG-CDX2 positive cells, suggesting OCT4 is essential for repression of *Cdx2* by NANOG (Figure 3D, S4D, S4E). Intriguingly CDX2 is expressed randomly in the *Nanog* null ESC colonies, unlike in WT where CDX2 was mostly restricted to the cells in the periphery of the ESC colony (Figure 3E). We analysed multiple ChIP-Seq data to identify possible regulatory elements for CHIR mediated induction of *Cdx2*. Intron-I region of *Cdx2* was enriched in OCT4, NANOG, TCF3 and β-CATENIN ChIP-seq data. Trophectoderm enhancer (TEE) region upstream of the promotor was enriched in YAP-TAZ ChIP-seq (Figure S4F). We cloned the *Cdx2* promotor (TSS), TEE, and part of intron-I; upstream of a minimal β-Globin promotor driving mCherry reporter (Figure 3F). Reporter cell lines were created by transgenic integration of these constructs in ESC. A significant induction of mCherry was observed in Intron-I reporter cell line after CHIR treatment suggesting CHIR might induce *Cdx2* through intron-I (Figure 3F). Further we deleted the TEE, Intestinal enhancer (IEE), OCT4 binding regions (OBR), NANOG binding regions (NBR) and both (ONBR) in TCMC. CHIR treatment failed to induce *Cdx2*:mCherry in ΔOBR, ΔNBR, and ΔONBR (*Cdx2*-Intron-I) confirming that *Cdx2*-Intron-I is essential for CHIR-mediated induction of *Cdx2* (Figure 3G). Inhibition of GSK3β by CHIR activates β-CATENIN, we asked if β-CATENIN is essential for induction of *Cdx2*. CHIR fails to induce *Cdx2*:mCherry in *β-Catenin* null TCMC (Figure S4G). Together our data suggest that CHIR acts through β-CATENIN on Intron-I of *Cdx2* to induce its expression. Both OCT4 and NANOG are together essential for repression for CHIR-induced expression of *Cdx2* and priming of ESC to TSCs.

**Fig. 3.**
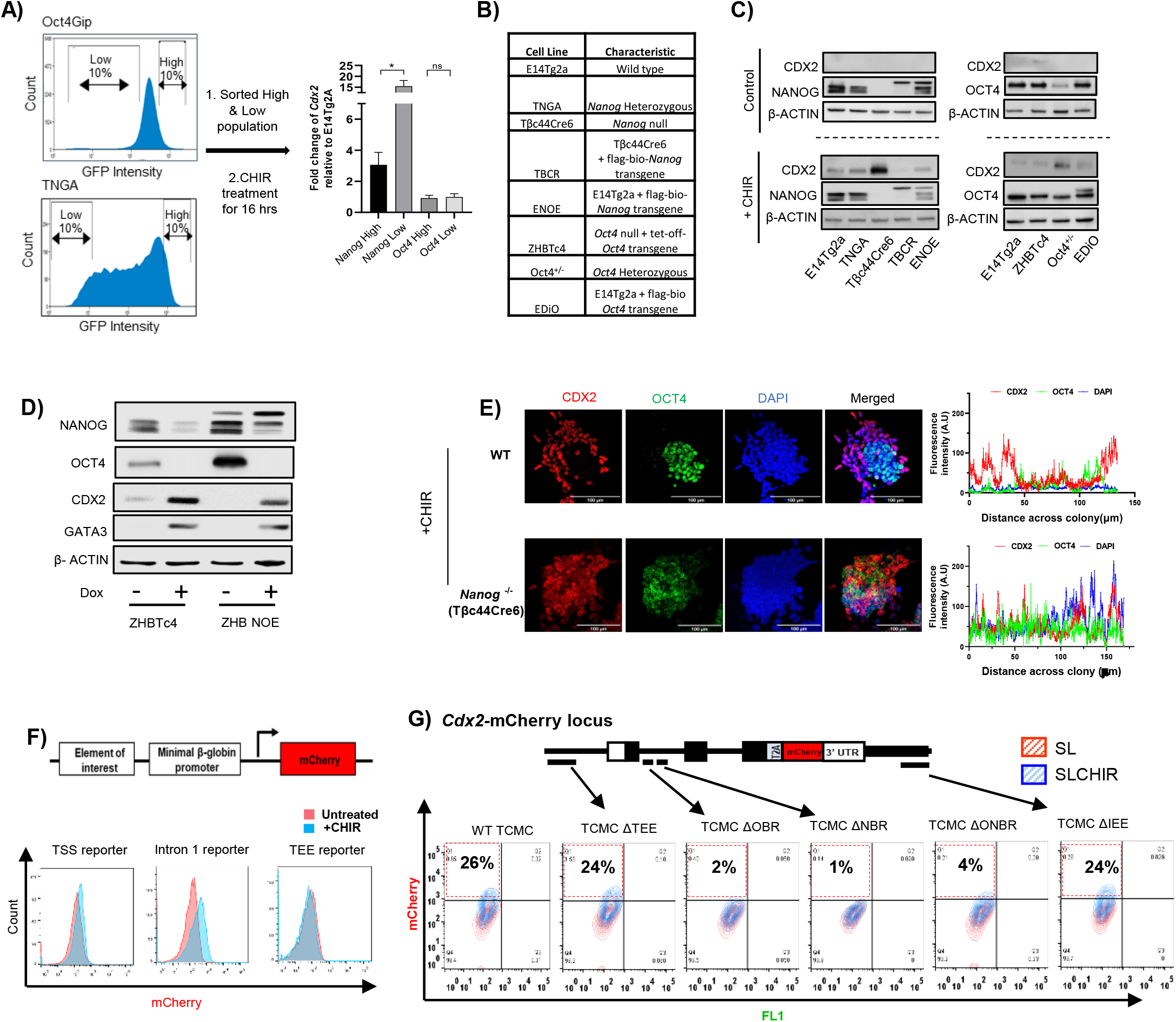
β-CATENIN and Intron-I regulatory region of Cdx2 are essential for priming of ESCs to TSCs. (Left) FACS profiles of TNGA and Oct4Gip cell line. The 10% low and 10% high GFP expressing cells were sorted, plated, and treated differentially with CHIR for 16 hours (Right) q-RTPCR analysis of Cdx2 expression relative to E14Tg2a in the 10% low and 10% high GFP expressing cells (n=3) ‘*’ indicates a p-value <0.05, ‘**’ indicates a p-value <0.01. B) List of cell lines describing the genetic manipulation of Oct4 and Nanog. C) (Left) Immunoblot analysis of CDX2 and NANOG in the indicated ESC lines cultured in SL and SLCHIR. (Right) Immunoblot analysis of CDX2 and OCT4 in the indicated ESC lines cultured in SL and SLCHIR. D) Immunoblot analysis of NANOG, OCT4, CDX2, GATA3, and β-ACTIN in ZHBTc4 (ZHB) and ZHBTc4 NOE (expressing a Nanog transgene). E) (Left) Immunofluorescence detection of CDX2 and OCT4 in CHIR-treated WT and Nanog null cell line. (Scale bar = 100µm). (Right) Distribution of fluorescence intensity of CDX2 and OCT4 across a colony. F) (Top) Schematic depicting the DNA constructs used for the mCherry reporter assay to assess the Cdx2 regulatory elements in ESCs. (Bottom) histogram profile of mCherry expression observed in stable ESC lines carrying Cdx2 regulatory elements reporter treated with or without CHIR. G) Contour plots showing the expression of mCherry in TCMC, TCMCΔTEE, TCMCΔOBR, TCMCNBR, TCMCΔONBR, and TCMCΔIEE treated with/without CHIR.

### β-CATENIN and Intron-I mediated activation of *Cdx2* is essential for localisation of ESC to the TE compartment of blastocyst

The ground state ESCs are cultured in 2iL(Ying et al., 2008), EPSCs (D-EPSCs from Deng laboratory) cultured in LCDM (LIF, CHIR, DiM, MiH)(Yang et al., 2017b), and Expanded PSCs (L-EPSCs from Liu laboratory) in EPSCM (CHIR, LIF, PD0325901, SB203580, A-419259, XAV-939)(Yang et al., 2017a). Intriguingly, apart from LIF, CHIR is the only common small molecule in all these culture conditions enabling enhanced pluripotency. We asked if CHIR-mediated induction of *Cdx2* through β-CATENIN and *Cdx2-* Intron-I is essential for the extraembryonic potential of the ground state ESC and EPSCs. *Cdx2*:mCherry expression was not induced in SLCHIR, 2iL, LCDM, and EPSCM in TCMC cells lacking β-CATENIN (TCMBβ-Cat-/-) implying an essential role for β-CATENIN in *Cdx2* induction (Figure 4A). *Cdx2*:mCherry was induced in TCMC cultured in SLCHIR, 2iL, and LCDM, but not in EPSCM (Figure 4B). Trophoblast factors CDX2 and GATA3 function downstream of TEAD4. Either factor can independently impart trophoblast fate when ectopically expressed in ESCs(Ralston et al., 2010). We asked which trophoblast factors are induced in the above stem cell states to induce trophoblast potential. CDX2 was detected in ESC cultured in SLCHIR, 2iL, and LCDM but not in EPSCM (Figure 4C, S5A). The absence of TEAD4 expression (Figure S5A) suggests that the induction of *Cdx2* in D-EPSCs is independent of TEAD4. We asked if *Cdx2-*intron-I is essential for *Cdx2* expression in these pluripotency states. *Cdx2*:mCherry expression was not induced in 2iL and LCDM in ΔOBR, ΔNBR, and ΔONBR suggesting that the intron-I regulatory regions of *Cdx2* are essential for the activation of *Cdx2* in 2iL (S5B). These data show that β-CATENIN and *Cdx2*-intron-I are essential for *Cdx2* induction in 2iL. However, *Cdx2* expression was not induced in EPSCM despite the presence of CHIR. We reasoned that the presence of XAV in EPSCM could increase stability of AXIN and might inhibit CHIR mediated activation of β-CATENIN and *Cdx2*. Addition of XAV in LCDM significantly reduced the activation of *Cdx2* confirming that XAV counteracts CHIR mediated induction of *Cdx2* (Figure 4B, 4C, S5A). Failure to derive ESTS from TCMBβ-Cat-/- and ΔONBR ESC suggest that the β-CATENIN and *Cdx2-*intron-I are essential for TSC-priming and derivation of ESTS (Figure 4D). Together our data demonstrate that CHIR is the key molecule in 2iL and LCDM that primes TE fate in pluripotent cells by induction of *Cdx2* through β-CATENIN and *Cdx2-*intron-I.

**Fig. 4.**
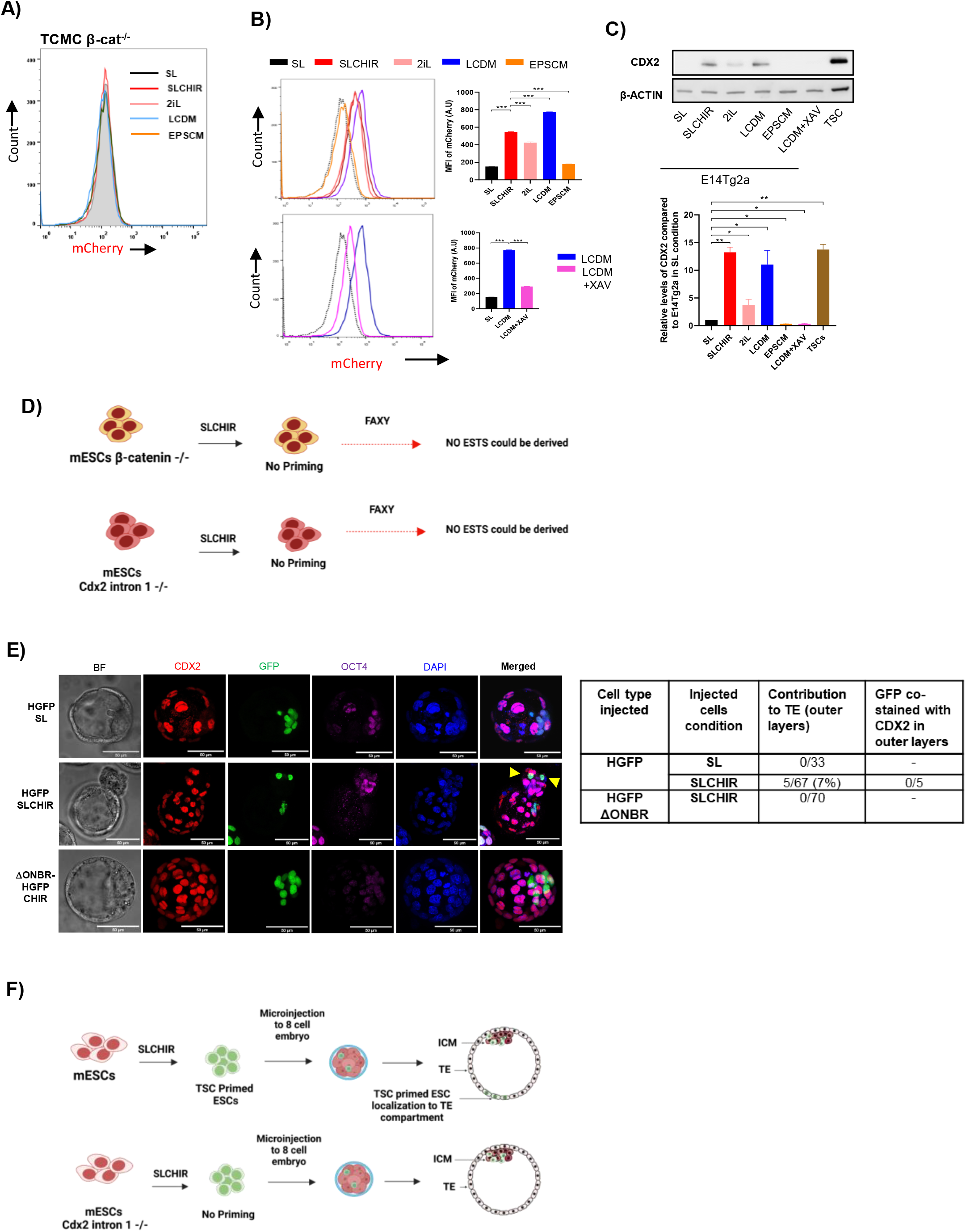
β-CATENIN and Intron-I mediated activation of Cdx2 is essential for localisation of ESCs and EPSCs to the TE compartment of the blastocyst. (A) mCherry expression in TCMCβ-Cat-/- cell line cultured in SL, SLCHIR, 2iL, LCDM, and EPSCM. (B) (Top left) mCherry expression in TCMC cell line cultured in SL, SLCHIR, LCDM, and EPSCM (top right) Relative quantification of mean fluorescence intensity of mCherry (n=3). (Bottom left) mCherry expression in TCMC cell line cultured in LCDM and LCDM+XAV. (Bottom right) relative quantification of mean fluorescence intensity of mCherry (n=3), ‘***’ indicates a p-value <0.001. (C) (Top) immunoblot analysis of CDX2 and β-ACTIN in E14Tg2a and TSC in indicated culture conditions. (Bottom) relative CDX2 levels as estimated by densitometry (n=3) ‘*’ indicates a p-value <0.05, ‘**’ indicates a p-value <0.01. (D) Schematic representation of method followed to derive ESTS from TCMCβ-Cat-/- and TCMCΔONBR cell lines. (E) (Left) Immunostaining of CDX2 and OCT4 in chimeric blastocysts developed from morula injected with HGFP and DONBR-HGFP ESCs cultured in SLCHIR or LCDM. (Scale bar =50µm). (Right) Table summarizing the outcomes of the chimeric blastocyst experiments. (F) Schematic representation of experiment to evaluate the chimeric potential of HGFP and DONBR-HGFP cell lines cultured in SLCHIR or LCDM.

We asked if CHIR-induced *Cdx2* could alone enable ESC to contribute to extraembryonic lineage *in vivo*. We injected H2B-GFP-ESC (HGFP) cultured in SL, and SLCHIR into morula (E2.5). The contribution of the injected cells to the three lineages of the blastocysts was analysed by immunostaining at E4.5. The descendant cells of HGFP cultured in SL contributed to the epiblast (Figure 4E). The cells cultured in SLCHIR (TSC-primed-ESC) were mostly localised to the epiblast. They also localised to the TE compartment in 7% of the blastocysts (Figure 4E) suggesting that ESCs cultured in SLCHIR could localize to TE compartment. The cells that localised to epiblast expressed OCT4, However, these cells in the TE compartment neither expressed OCT4 nor CDX2 (Figure 4E). This strengthens our findings that induction of *Cdx2* by CHIR primes the ESC to TE lineage and such priming may be sufficient for their localization to TE compartment. However, the absence of CDX2 expression in derivatives of TSC-primed-ESC suggest that these cells do not acquire bona fide TSC capability. They need additional coaxing cues for the complete transition to TE lineage. Our data is similar to the lack of CDX2 and OCT4 expression in D-EPSCs localised to the TE position as reported by Posfai et. al.(Posfai et al., 2021).

We asked if the ability of the ESC in SLCHIR to localise to the TE position in blastocysts is merely due an intermediate stage resulting from induction of *Cdx2* by CHIR or an overall extended potential of the cells. We generated TCMC-ΔONBR ESC carrying an H2B-GFP transgene (ΔONBR-HGFP). The cells were cultured in SLCHIR and injected into morula stage embryos to analyse theirs *in vivo* developmental potential. The injected cells exclusively contributed to the epiblast and failed to localize to TE position, suggesting *Cdx2*-Intron-I is essential for the localisation of these cells to the TE position in the blastocyst (Figure 4E, 4F). Our data suggest that the ability of the TSC-primed-ESCs to localise to TE position in developing blastocyst can be attributed to *Cdx2* induction through Intron-I and may not be due to the overall extended potential of these cells.

### Derivation of TSCs from mouse epiblast

The ability to develop into the TE lineage is unconstrained in human epiblast but is considered to be lacking in mouse epiblast(Guo et al., 2021). ESTS can be derived from mESC in FAXY, we investigated whether the mouse epiblast had the same potential to differentiate to TE linage. The plasticity of mouse ICM to form trophoblast is lost by mid blastocyst stage(Posfai et al., 2017). We isolated ICM of the mouse embryo after E3.75 by immunosurgery(Solter and Knowles, 1975) and cultured in SL and SLCHIR for 24 hrs. The media was changed to FAXY and cultured for one week (Figure 5A). Outgrowths appeared in both culture conditions. Surprisingly the cells of the outgrowth obtained from SLCHIR→FAXY morphologically resembled endoderm, whereas some of the outgrowths from SL→FAXY resembled TSC morphology (Figure 5B). The outgrowths were picked and cultured further in FAXY. TSC-like cell lines (EpiTS) could be established from the SL→FAXY epiblasts. The EpiTS expressed CDX2 and could self-renew for multiple passages (Figure 5C). The possibility of residual TE cells surviving the immunosurgery giving rise to TSCs cannot be ruled out. To evaluate this possibility, we performed immunosurgery on 96 blastocysts. Half of the embryos (48 nos) were randomly chosen for immunostaining with CDX2 and OCT4. The other half (48 nos) were cultured in SL→FAXY condition. Outgrowths from 8 out of 48 (16%) epiblasts could be established into EpiTS. Only one cell in the epiblast of an embryo (2%) was observed to stain for CDX2 after immunosurgery (Figure 5D, 5E) suggesting only 2% of epiblast contained contaminating CDX2 expressing cell. This suggests that the TSC outgrowth appearing after immunosurgery arise mostly from epiblast. To scrutinize this further, we utilized HGFP cells which can contribute only to ICM of the developing blastocyst (Figure 4E) when cultured in SL. We injected HGFP cells cultured in SL into 8 cell embryos and subjected the chimeric blastocyst to immunosurgery and EpiTS derivation (Figure 5F). The immunostaining of the outgrowths from the HGFP chimeric epiblast showed some of the cells co-expressing GFP and CDX2 (Figure 5G) suggesting the epiblast cells were indeed differentiating to TE lineage. The GFP-CDX2 co-expressing cells were found in 20% of the outgrowths (Figure 5G). Collectively, we show that mouse epiblast possesses potential to differentiate to TE linage and EpiTS can be derived from mouse epiblast.

**Fig. 5.**
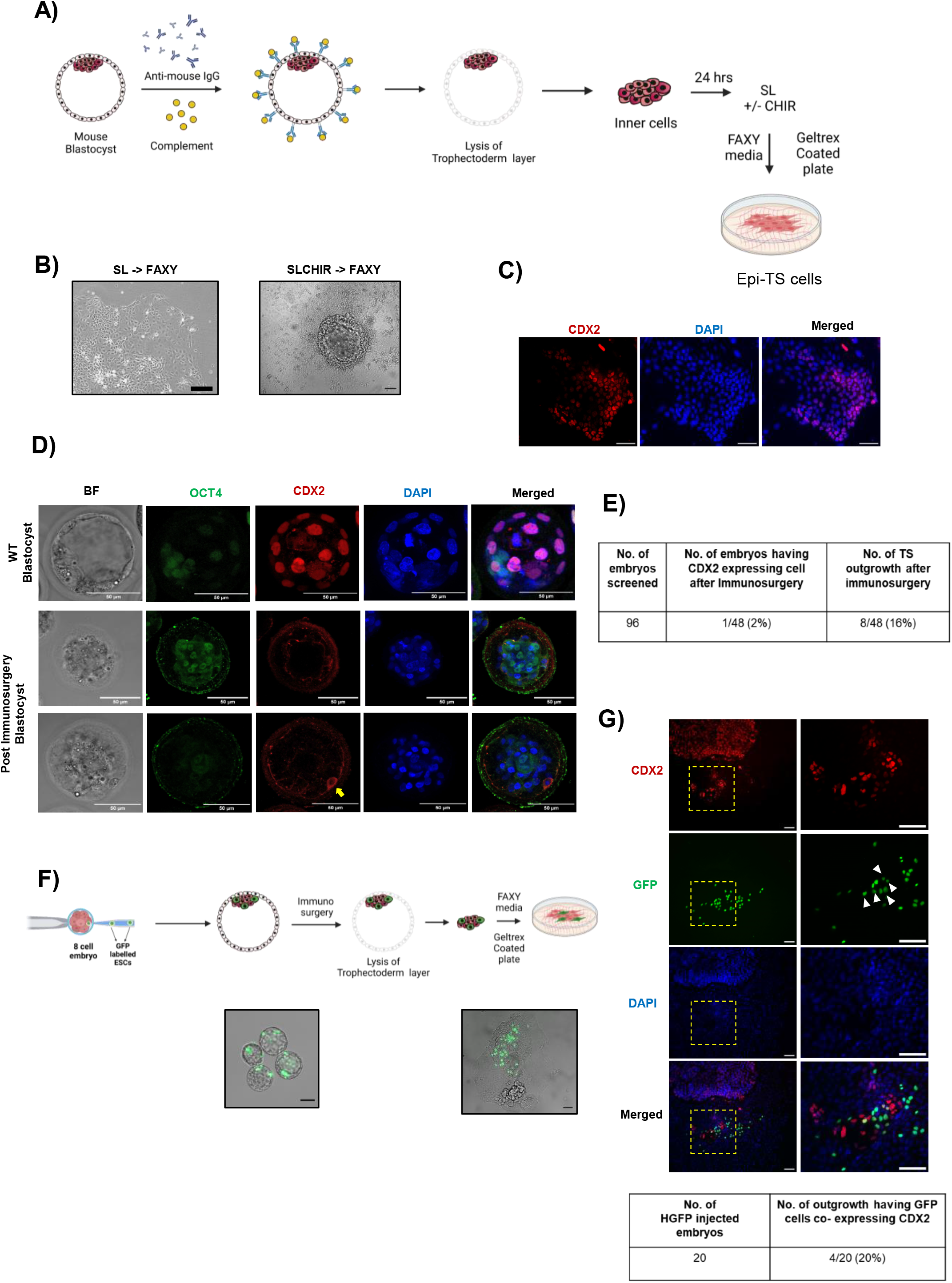
Derivation of TSCs from mouse epiblast. A) Schematic of the derivation of TS-like cells from E3.75 mouse blastocyst (EpiTS). B) Phase contrast image of TS-like outgrowth cultured from post-immunosurgery blastocyst under depicted culture conditions. C) Immunostaining of CDX2 in EpiTS derived from ICM. Scale bar = 50µm. D) Immunostaining of CDX2 and OCT4 in embryos following immunosurgery treatment. (Top row) Immunostaining E3.5 blastocyst (Middle and bottom row) Immunostaining of E3.5 embryo post immunosurgery. The yellow marked arrow marks the single CDX2 positive cell in an embryo out of 48 embryos analysed. Scale bar = 50µm. E) Table summarizing the number of embryos having CDX2 expressing cells and number of TS-like outgrowth after immunosurgery. F) (Top) Schematic showing injection of GFP labelled ESCs (HGFP) into 8-cell stage embryo followed by EpiTS derivation from the chimeric blastocyst. (Left Bottom) Phase contrast image overlaid in GFP channel showing the contribution of HGFP in E3.5 blastocyst. (Right Bottom) The Phase contrast image overlaid in the GFP channel shows the contribution of the HGFP Epi-TS colony. Scale bar = 50µm. G) (Top) Immunostaining of CDX2 and GFP in EpiTS colony. (Bottom) Table summarizing the number of HGFP-injected embryos and the number of TS-like outgrowth coexpressing GFP and CDX2. Scale bar = 50µm.

## Discussion

In this study, we have derived TSC from murine ESC by a two-step process. First involving a TSC-priming of ESCs to activate the pioneer transcription factor *Cdx2*(Liang et al., 2021) followed by a complete transition to TSC. *Cdx2* transgene expression can convert ESCs to TSCs(Kuckenberg et al., 2010; Niwa et al., 2005). We show that GSK3β inhibition induces *Cdx2* expression in ESCs, however such cells fail to convert to TSC in presence of a GSK3β inhibitor. This could be attributed to pleiotropic effects such as the upregulation of *Stat3* by GSK3β inhibition(Hao et al., 2006), which might interfere in the next stage of complete TSC transition. The essential function of XAV in the second stage of complete TSC transition and failure to derive ES-TSC in the continued presence of CHIR in FAXY supports a stage-specific function for GSK3β in ESTS derivation. Inability of the other TSC media(Okae et al., 2018; Tanaka, 2006) to enable the complete TSC transition underscores the essential stage-specific role of Gsk3β function in ESTS derivation. Expression of *Gata3*, *Tead4*, *Raf*, *Eomes*, and *Tfap2c* can induce TE lineage genes similar to *Cdx2* in ESCs or fibroblasts to generate TSCs(Ng et al., 2008; Nishioka et al., 2009; Niwa et al., 2005; Ralston et al., 2010). We suggest, it might be possible to develop novel culture conditions to derive TSCs from PSCs following a similar approach of TSC-priming of ESCs by small molecule induction of any of these factors or their combinations.

The TSC-priming step induces some TE lineage markers including *Cdx2* in ESCs. Although such cells do not acquire bonafide TE lineage potential, they can still localise to the TE compartment in balstocysts. *Cdx2*-intron-I dependent activation of *Cdx2* is essential for TSC-priming and TE compartment localisation of ESCs and EPSCs. We suggest that the rare sporadic contribution of ESCs to the TE compartment in embryos observed by us and others(Beddington and Robertson, 1989; Posfai et al., 2021; Yang et al., 2017b) is a result of the localisation of descendants of TSC-primed pluripotent cells which lack bonafide TE potential.

Unlike the human epiblast, mouse epiblast and ESCs are considered to lack the potential to readily contribute to trophoectoderm lineage(Beddington and Robertson, 1989; Gardner, 1983; Guo et al., 2021; Nichols and Gardner, 1984; Posfai et al., 2017). The chromatin-based lineage barriers were speculated to restrict the Trophoectoderm potential of the mouse epiblast and ESCs(Boyer et al., 2006; Ng et al., 2008; Zijlmans et al., 2022). Recently the histone posttranslational modifications (hPTMS) were found to be largely persevered between hPSC and mESCs suggesting, chromatin barriers may not be major mechanism by which the mouse epiblast/mESC potential is restricted(Zijlmans et al., 2022). Failure to derive TSC from mouse ESC by us and others using the methods used for human TSC and derivation of ESTS in FAXY reaffirms this proposition.

The extra embryonic lineages subserve the development of the epiblast to ensure successful development of the progeny and perpetuation of the species. The segregation of epiblast and extraembryonic layers are essential for proper compartmentalization and development. Any damage to the extraembryonic layer cells can affect the development of the epiblast. Hence, we propose that the mammalian epiblast has inherent plasticity to differentiate to extraembryonic layers well beyond the timelines of these linage segregations during development. This inherent potential is restricted by multiple developmental signals and the organization of the embryonic layers. In instances of damage to the extra embryonic layers the epiblast cells can differentiate to the cells of TE or PE compartment to compensate for the lost cells and ensure continued development of the epiblast. The lineage restriction imposed by the developmental signals on the epiblast may be overcome by the signals generated by the damage to the extraembryonic layers. This is best exemplified from our immunosurgery experiments and by Guo et. al., where the damage to TE leads to differentiation of TE from mouse and human epiblast and further supported by development of reformed blastocoel from the ICM of bovine with a TE layer. The development of blastoids entirely from PSCs of human(Kagawa et al., 2022), monkey(Li et al., 2023) and mice(Jana et al., 2023) supports the existence of inherent plasticity in mammalian epiblast and PSCs to generate all extraembryonic lineages.

Collectively, our data demonstrate that the mouse ESCs have unrestricted developmental potential contrary to the prevailing understanding. Under appropriate conditions, the unrestrained potential of mESC can be unlocked to derive TSCs. The demonstration of the unrestrained developmental potential of murine and human PSCs opens up the possibility to harness the extraembryonic and blastoid generation potential in other mammalian PSCs by developing appropriate culture approaches.

We suggest that there are multiple layers of lineage barriers -like signaling barriers (mechanical and cytokine), Pluripotency factors, chromatin-based barriers operating in different mammalian species which need to be overcome for realization of the full developmental potential of the epiblast. We speculate that a typical mammalian epiblast is not governed by strict adherence to restricted lineage but by an adaptive lineage potential which restrict TE line under normal development, but can permit differentiation to TE linage when the need arises.

## Acknowledgments

D.J, V.V.V, M.S was supported by a fellowship from UGC (India). H.T.K was supported by a fellowship from ICMR (India). We thank the Microscopy, FACS, Animal House and transgenic core facilities of CCMB for the support extended to carry out this work. P.Si acknowledges the stipend support from the DBT grant GAP0546. WT/DBT India Alliance grant 500053/Z/09/Z -P.C.S., Department of Biotechnology grant BT/PR14064/GET/119/16/2015-P.C.S., DBT grant GAP0582: BT/PR40264/BTIS/137/44/2022 -D.T.S.

## Author contributions

Conceptualization, D.J, and P.C.S Methodology, D.J, P.Si, P.Sa, J.L, G.S, and P.C.S; Investigation, D.J, P.Si, P.Sa, and D.T.S; Writing – Original Draft, D.J, P.Si and P.C.S.; Writing – Review & Editing, D.J, P.Si, D.T.S and P.C.S.; Funding Acquisition, D.T.S and P.C.S.; Resources, D.J, M.S, V.V.V, and H.T.K; Visualization, P.Si and D.J.; Supervision, P.C.S.

## Methods

### Culture of metastable and ground-state mouse ES cells

Metastable ES cells were grown on 0.1% (w/v) gelatin-coated cell culture dishes in serum+ LIF media, comprising 10% (v/v) heat-inactivated FBS in GMEM (12.5g/litre w/v), NaHCO3 (32.7mM), Sodium Pyruvate (1mM), NEAA (0.1mM) and β-mercaptoethanol (0.1mM) and LIF. Ground state ES cells were cultured on 0.1% gelatine-coated tissue culture-treated dishes in N2B27, containing DMEM/F12 and Neurobasal medium in 1:1 (v/v) ratio, supplemented with N2 supplement (1X), B27 supplement (1X), NaHCO3 (32.7mM), Sodium Pyruvate (1mM), NEAA (0.1mM), and β- mercaptoethanol (0.1mM) along with recombinant hLIF (1000U/ml), CHIR99021 (3μM) and PD0325901 (1μM). TrypLE-EDTA was used for dissociation and passing once the confluence reached 70%.

### Extended pluripotent stem cell culture

Metastable ES cells were converted into Extended pluripotent stem cells (EPSCs) when grown in either of two different culture regimes. First as described by Pentao Liu’s group(Yang et al., 2017b), where DMEM/F12 (Invitrogen), 20% (v/v) KnockOut Serum Replacement (KSR), NEAA (1X), β-mercaptoethanol (0.1mM) and hLIF 1000 U/ml supplemented with the following small-molecule inhibitors: CHIR99021 (3μM), PD0325901 (1μM), JNK Inhibitor VIII (4μM), SB203580 (10μM), A-419259 (0.3μM) and XAV939 (5μM). The second culture media as described by Hongkui Deng’s group(Yang et al., 2017b) where basal N2B27 media was supplemented with small molecules and cytokines as follows: 10 ng/ml recombinant hLIF, CHIR99021 (3µM), (S)-(+)-Dimethindenemaleate (2µM), Minocycline hydrochloride (2µM) and 5mg/ml BSA. Upon 70% confluency, EPSCs were passaged with TrypE-EDTA, and splitting was done in a 1:3 ratio.

### Derivation of ESTS from ES cells and TSC from blastocyst

For the derivation of trophoblast stem cells (ESTS) from mESCs, E14Tg2a cells were cultured for 48hrs in SL media supplemented with CHIR99021 (3μM). Thereafter, cells were passaged and seeded onto culture plates as described by Ohinata et. al.(Ohinata and Tsukiyama, 2014). Geltrex (1x) or 15µg/ml fibronectin solution in PBS was used to coat the tissue culture-treated plates at 37oC for 1 hour. SLCHIR-treated E14Tg2a cells were seeded onto these plates and cultured in CDM/FAXY media. CDM/FAXY media contained DMEM/F12: Neurobasal media (1:1 v/v), supplemented with N2 (1X) and B27 (1X) supplements, BSA (0.05% v/v), 1-thioglycerol (1.5 X 10-4M), recombinant human bFGF (25ng/ml), recombinant human Activin A (20 ng/ml), XAV939 (10 nM), and Y27632 (5nM). By day 6, colonies with TS-like morphology were surrounded by contaminating non-TS-like cells. Two methods were used to separate ESTS from the contaminating cells. First, by manually picking the colonies having TS-like morphology with a fine pipette and seeding them back into coated dish with FAXY media as a single cell suspension. In another method, CDM/FAXY media was removed, and the culture was treated with Accutase for 30sec or until the contaminating cells started to detach from the dish after gentle tapping. The dishes were washed with PBS to remove the detached cells, while the TS-like cells remained attached. This method utilises the deferential sensitivity of the TS-like cells and the contaminating cells to the Accutase treatment. Media was changed every 24hrs and clearing of contaminating cells was done after the cell density reached 60% confluency. The first cleaned ES-TSC plate was named ESTSP1 followed by multiple passages as P2-P24. ESTSP24 or later passages are hereinafter called ESTS. TSC were derived from blastocysts as described in Ohinata et. al.(Ohinata and Tsukiyama, 2014).

### Mice

CD1 and F1 (C57BL6/CBA) mice were obtained from Jackson laboratories. Animal model experiments were carried out following the Institutional Animal Ethics Committee (IAEC) of the CCMB’s ethical standards and received approval from the Committee for Control and Supervision of Research on Animals (CPCSEA). For pseudo-pregnancy, Vasectomized F1 male mice were mated with CD1 female mice that were 8 to 11 weeks old and in the estrus cycle. All the experimental mice were kept in a 12-hour light/12-hour dark cycle with free access to food and water in a facility with controlled temperatures (18-22°C) and humidity (R.H. 40-70%).

### Superovulation of mice

5 IU of PMSG (pregnant mares’ serum gonadotrophin) was administered to female F1 (C57BL6/CBA) for superovulation. 5 IU of human chorionic gonadotrophin (HCG) was administered 48 hours after the initial injection and Day 0 was designated of and the subsequent instant mating with F1 male mice. After the vaginal plug was found the following day, 0.5 dpc zygotes were retrieved and cultivated in KSOM in an incubator that was humidified and maintained at 37°C.

### Culture of mouse embryos

The upper portion of the oviduct (ampulla) was cut-off from the pregnant mice, and zygotes (0.5dpc) and cumulus mass were extracted from F1 (C57BL/6J X CBA). A prepared M2 medium with 0.3 mg/ml hyaluronidase was used to transfer the oviduct area containing the zygotes. Close to the location of the zygotes, the ampulla was torn using a 26-gauge needle. For a short period of time, all of the zygotes were incubated in M2 medium containing hyaluronidase. Zygotes were picked up using a pipette after the cumulus mass had fallen off and transferred to new M2 drops before being recorded as Day 1 and pre-equilibrated at 37°C with 5% CO_2_ in a humid incubator. Every 24 hours, embryos were transferred into fresh KSOM drops.

### Immunosurgery of E3.5 blastocyst

The immunosurgery technique was used to separate the inner cell mass (ICM) and trophectoderm (TE) of mice E3.5 blastocyst(Solter and Knowles, 1975). The blastocysts were transferred to the M2 medium and washed twice in the M2 medium. Blastocysts were incubated with Rabbit anti-mouse antibody at a dilution of 1:10 (diluted in DMEM plus 10% FBS) for 15 minutes at 37°C in a CO2 incubator. Blastocysts were washed three times with M2 medium after the incubation. Blastocysts were transferred to a fresh drop containing native guinea pig serum complement at a dilution of 1:50 (diluted in DMEM plus 10% FBS), and incubated for 30 minutes at 37°C in a CO2 incubator. The blastocysts were then observed under a microscope to confirm the lysis of the TE cells. Zona pellucida was cleared with acid Tyrode treatment for 2 mins. The ICMs were collected carefully using a micropipette and transferred to a new dish with fresh M2 medium. Finally, the ICMs recovered were transferred to Geltrex coated cell culture plate containing FAXY media for EpiTS differentiation.

### Injection of ESCs, CHIR-treated ESCs, and ESTS into an 8-cell embryo

GFP expressing ES (HGFP) (cell line pedigree chart in supplementary data) cells were cultured on SL, SLCHIR (CHIR, 3µM) for 16hrs. SLCHIR media was changed freshly 3hrs before injection. For ESTS transfer, GFP labelled ESTS cells which were grown in FAXY for 24 passages (ESTSP24) or later passages cells were trypsinized with TrypLE-EDTA, neutralized, washed, and resuspended as single cells in injection media. Injection media was freshly made and constituted of DMEM/F12 (Sigma) supplemented with 10% (v/v) heat-inactivated ES qualified serum, Glutamax (1X), NaHCO3 (32.7mM), and pH was adjusted to 7.2. Resuspended ESCs were microinjected into the 8-cell stage embryo in an injection drop containing 200µl of injection media in a 50mm glass-bottom dish. Leica DM IRB inverted microscope was used with an Eppendorf TransferMan NK2 micromanipulator. The outside and inside diameters of the holding pipette were 95–105μm and 20–25μm respectively; whereas the outside and inside diameters of the injection pipette were 18–20μm and 16–18μm respectively. 2-3 ESCs/ESTS cells were microinjected per embryo on a 50mm dish containing injection media on a microscopic stage maintained at 100C. A batch of 20 embryos was injected in an hour.

### Cryosectioning of placental tissue and cytohistochemistry

Placental tissues were obtained after dissecting 12.5 dpc mice. Tissues were immediately fixed in 4% paraformaldehyde and kept overnight at 4°C. Tissues were washed thrice with PBS and transferred to 15% sucrose solution in PBS and kept at 4°C until the tissues were sunk at the bottom of the tube. The tissues were again transferred to 30% sucrose solution in PBS and kept at 4°C overnight followed by OCT media embedding and taken for cryosectioning. Sections of 15-micron thickness were taken on ProbeOn slides and followed for immunocytochemistry. The slides were dried at room temperature for 5mins and washed with PBS twice to remove the OCT. Sections were blocked and permeabilized in a blocking buffer containing PBS with 5% (w/v) BSA, and 0.3% (v/v) Triton-X 100 for 60 minutes at room temperature. Primary Antibodies were diluted in antibody dilution buffer (ADB) containing PBS with 1% (w/v) BSA, and 0.3% (v/v) Triton-X 100. Sections were incubated with primary antibodies for 4°C overnight. Sections were washed 3 times with PBS and incubated with secondary antibodies diluted at 1:1000 in ADB. Sections were washed with PBS and mounted with Vectashield (H-1200) containing DAPI.

### Generation of CRISPR-based gene knock-out cell lines

The twin guide strategy was used for generating knock-out in ES cells as described in Kale et. al.(Kale et al., 2022). The cell line was characterized by genotyping, qPCR, and western blotting analysis to confirm the intended genetic manipulation. The detailed characterization and pedigree of the cell lines used in this study are provided in supplementary data (supplementary appendix).

### Generation of ES cells knock-in cell lines

TCMC cell line was generated by using CRISPR as described by Jana et al. (Jana, et al. 2019). TCMC-OGFP cell line was generated by nucleofecting 1µg supercoiled Oct4-IRES-eGFP-PGK-Neo (Addgene #48681) plasmid to 1 million TCMC using P3 primary cell 4D-Nucelofector X kit (Lonza). Nucleofected cells were plated and grown in SL media supplemented with G418 for 10 days. A single colony was picked up and cultured as replicas in 96 well format for genotyping. The detailed characterization and pedigree of the cell lines used in this study are provided in supplementary data (supplementary appendix).

### Immunofluorescence staining

Cells were cultured in 2D culture in 24 wells for up to 70% confluency. Cells were washed thrice with 500µl of PBS and 500 µl of freshly prepared 4% paraformaldehyde (made in PBS) fixative was added to the plate and incubated at RT for 20 minutes. The fixative was removed and the plate was washed thrice with 1ml PBS. The specimen was blocked and permeabilized in a blocking buffer containing PBS with 5% (w/v) BSA, and 0.3% (v/v) Triton-X 100 for 60 minutes at room temperature. Primary Antibodies were diluted in antibody dilution buffer (ADB) containing PBS with 1% (w/v) BSA, and 0.3% (v/v) Triton-X100. Specimens were incubated with primary antibodies for 4°C overnight. Cells were washed 3 times with PBS and incubated with secondary antibodies diluted at 1:1000 in ADB. Cells were washed with PBS and mounted with Vectashield (H-1200) containing DAPI.

For staining 3D structures like embryos, they were first washed twice with PBS. Structures were fixed in 4% paraformaldehyde in PBS for 15 minutes and rinsed in PBS containing 3 mg/ml polyvinylpyrrolidone (PBS/PVP). Thereafter structures were permeabilised in PBS/PVP containing 0.25% Triton X-100 for 30 minutes. Blocked in blocking buffer, comprising PBS containing 5% BSA, 0.01% Tween 20 for 60 minutes. Primary antibodies were diluted with the appropriate antibody dilution as per the manufacturer protocol in PBS containing 1% BSA, and 0.01% Tween 20 and incubated at 4°C overnight. They were rinsed three times in blocking buffer for 5 minutes each and incubated with secondary antibodies diluted as 1:500 in PBS containing 1% BSA, 0.01% Tween 20, and incubated for 60 minutes at room temperature. Rinsed 3 times with PBS and stained for nuclei with DAPI (1µg/ml) prepared in PBS for 15 minutes at RT. Embryos were finally rinsed 3 times in PBS and images were acquired by confocal microscopy.

### Real-time PCR analysis

The RNA was extracted from 1 million cells with TRIzol by manual method and quantified by Nanodrop (Thermo Scientific). One microgram of RNA was reversed and transcribed into cDNA by using SuperScript™ III First-Strand Synthesis System. The first strand synthesized cDNA was diluted 5 times and real-time PCR was set with power SYBR Green PCR master mix on the ABI 7900 HT. The PCR setup was as follows: Step 1: 95°C for 10 min, step 2: 95°C for 15 sec, step 3: 60°C for 30 sec, and Step 4: 72°C for 30-sec Steps 2-4 were repeated for 40 cycles. GAPDH was used as an internal control. The reactions were analyzed by the software (SDS 2.1) provided with the instrument. The primers used for real-time PCR are given in the resource table.

### Western blot analysis

The cells were harvested by scraping them from the plates in PBS and collected by centrifugation. The cell pellet was washed twice with PBS and reconstituted in RIPA. RIPA buffer constituting 25mM Tris HCl (pH 8.0), 150mM NaCl, 1% NP-40, 0.5% Sodium deoxycholate, 0.1% SDS, and Complete Protease Inhibitor Cocktail Tablets (Roche). The lysate was sonicated in Bioruptor (Diagenode) and centrifuged. The clear supernatant was transferred to the fresh tube. The protein samples were denatured in loading dye containing β-mercaptoethanol and resolved by SDS-PAGE. Resolved samples were then transferred onto a polyvinylidene difluoride (PVDF) membrane and blocked-in blocking buffer containing 5% non-fat milk in TBST. Primary antibody hybridisation was carried out overnight at 4°C in 3% non-fat milk in TBST. After incubation three washes were performed with TBST. The secondary antibody was diluted at 1:10,000 in 3% non-fat milk in TBST and incubated for an hour at RT. After incubation three washes were performed with TBST and visualized using enhanced chemiluminescence (ECL) detection kit (Thermo Scientific) and developed in Chemi doc MP (Biorad).

### FACS analysis

70% confluent culture was made into single-cell suspension using TrypLE for 4 mins at 37°C. The cells were diluted in media and pelleted by spinning at 300g for 5 mins. The media was removed and around 1 million cells were resuspended in 300 µl of PBS containing 2% FBS. The cell samples were directly taken for analysis either in Gallios (Beckman Coulter B5-R1-V2) FACS analyzer or LSR Fortessa (BD) analyzer and data was recorded. The FACS data were analyzed using FlowJo vX.0.7

### Bulk RNA-seq library preparation

Using the TRIzol reagent and the manufacturer’s instructions, total RNA was extracted from ESCs, TSCs and ESTSCs. A total RNA of 1μg was used to generate libraries using of Illumina stranded total RNA prep with Ribo-Zero Plus library preparation kit (Illumina, 20040529). The Qubit dsDNA HS (High Sensitivity) Assay Kit (Invitrogen, Q32854) was used to measure library concentration, and different libraries were combined to create the final pool at an equimolar ratio. Using an Illumina NovaSeq 6000 instrument, libraries were sequenced for a read length of 151 bp reads and a read depth of approximately 30 million reads.

### Bulk RNA-seq data analysis

Paired-end Bulk RNA sequencing was done for ESCs, TSCs and ESTSCs. The quality assessment of raw data was done by FastQC v0.11.9 and Illumina universal adapter content was removed using cutadapt v2.8. The filtered sequencing reads were mapped to the mouse reference genome mm10 and Gene counts were obtained by STAR_2.5.4b. The data showed an average of 81% uniquely mapped reads which were annotated with the Ensembl database. Transcripts Per Kilobase Million (TPKM) were calculated by the rsem-calculate-expression function of RSEM v1.3.3. DESeq2 v1.30.1 was used to get the differentially expressed genes based on the Bayes theorem. Genes showing expression of at least 50 in a row were retained for further analysis. Principal component analysis (PCA) and heatmap analysis were performed with the functions plotPCA and pheatmap in R. The visualization of differential expression of marker genes was performed on TPM counts after scaling and normalizing the read counts by row.

### Single RNA-seq library preparation

Single-cell suspension was made from ESCs, TSCs and ESTSCs using the TrypLE-EDTA solution. Live cells were sorted as PI-negative populations using a FACS sorter. Approximately 5000 cells were loaded for Library preparation using Chromium Next GEM Single Cell 3’ Reagent Kits v3.1 (10x genomics). Libraries were sequenced for a read length of 91 bp reads.

### Single-cell RNA-seq data processing

Single-cell RNA-sequencing was performed using the 10X Genomics Chromium system and Illumina base call files (BCL) were demultiplexed using the mkfastq function of cellranger v6.0.2 which is specific to 10X libraries. Reads were aligned against mouse reference mm10 and filtered by the count pipeline to get the feature, barcode, and gene matrices. The R package Seurat v4.0.1 was used to analyze the feature-barcode matrix. The quality assessment of data was done by scater v1.18.6. Scrublet, a python tool, was used to calculate doublet scores and predictions. Cells with more than 500 detectable genes with a doublet score of <0.25 and expression of mitochondrial genes accounted for less than 5% of total expression were filtered from the dataset for further downstream analysis. The final dataset includes ESC, ESTS, and TSC derived from blastocysts. Samples were integrated using the merge function in the Seurat suite. Normalization and variance stabilization was done by sctransform and variations due to genes were regressed out. Cells were clustered with the FindClusters function with a resolution of 0.5 and visualization was done using the RunUMAP function.

### Data and materials availability

All the cell lines and plasmid constructs used in this study will be made available against and email request and Material Transfer Agreement (MTA). Raw and processed transcriptome data is deposited in NCBI GEO and is available under the accession GSE219001. The code used for analysis and visualization of scRNAseq data is available at https://github.com/SowpatiLab/ES-TSCs

### Statistical analysis

Statistical analysis was done by using a two-tailed paired student t-test. The representation of data is in the form of means+/-SDM. The was calculated for more than three independent experiments P value<0.05 is considered statistically significant. * represents P<0.05, ** represents P<0.01, *** represents P<0.001, and **** represents P<0.0001.

### Supplementary Figures

**Supplementary Fig. 1.**
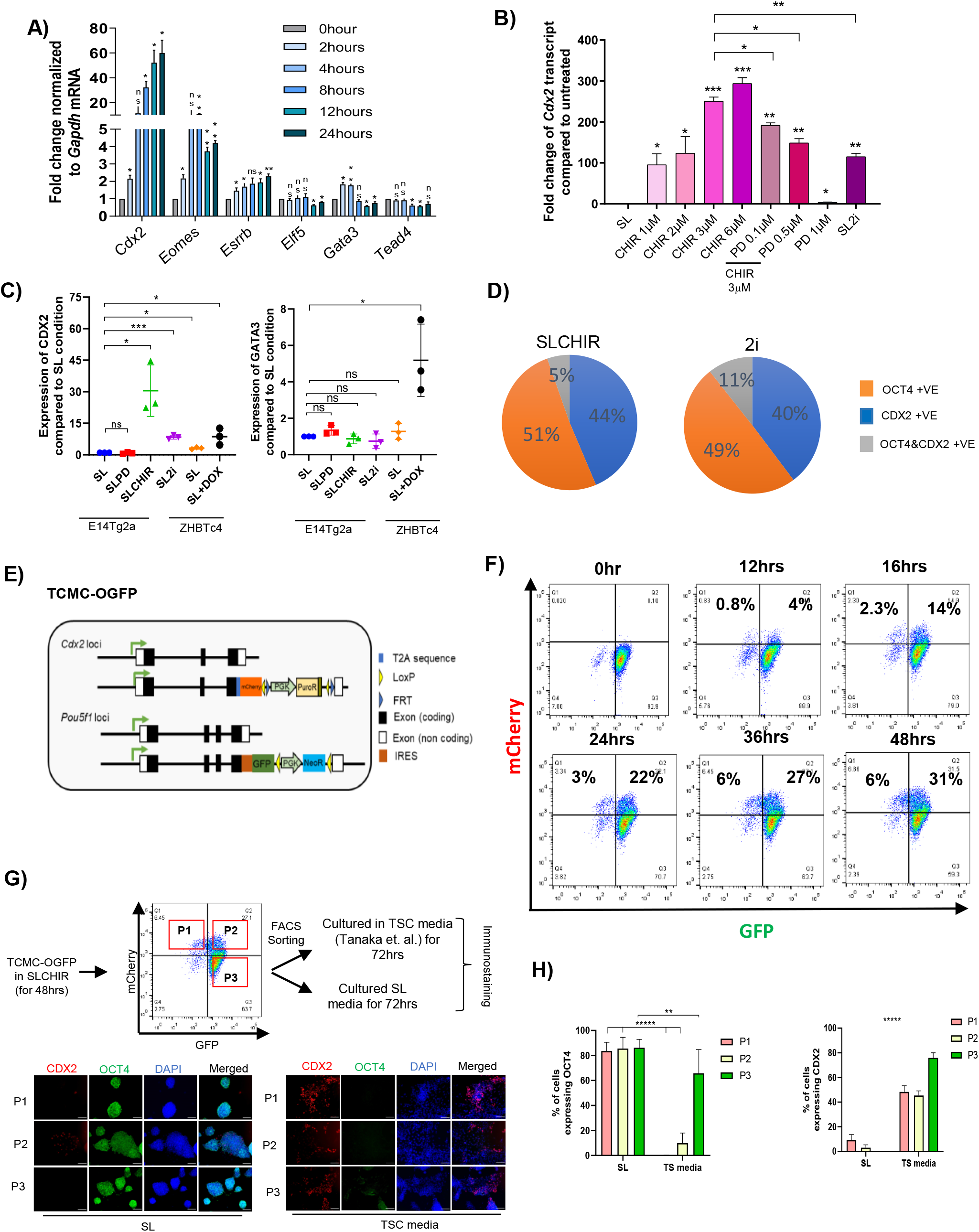
(A) q-RTPCR analysis of expression of TE lineage genes at indicated timelines after CHIR treatment. error bar represents the SD of the mean of biological replicates (n=3). ‘*’ indicates a p-value <0.05, ‘**’ indicates a p-value <0.01, ‘ns’ indicates a p-value > 0.05. (B) Expression of Cdx2 transcript in ESCs cultured in indicated combinations and concentration of PD and CHIR relative to ESCs cultured in SL. The error bar represents the SD of the mean of biological replicates (n=3). ‘*’ indicates a p-value <0.05, ‘**’ indicates a p-value <0.01, ‘***’ indicates a p-value <0.001, ‘ns’ indicates a p-value > 0.05. (C) Relative CDX2 and GATA3 levels as estimated by densitometry. The error bar represents the SD of the mean of biological replicates (n=3). ‘*’ indicates a p-value <0.05, ‘**’ indicates a p-value <0.01, ‘***’ indicates a p-value <0.001, ‘ns’ indicates a p-value > 0.05. (D) Pie chart indicating the fraction of cells expressing OCT4 or CDX2 or both analyzed from the immunofluorescence data. (E) Schematic depiction of TCMC-OGFP ESC line reporting Cdx2 and Oct4 expression by mCherry and GFP respectively. A T2A-mCherry cassette is integrated in-frame with the last coding sequence of one allele of the Cdx2 gene. An IRES-GFP cassette is integrated into the 3’UTR of one of the alleles of the Oct4 gene. (F) FACS profile for TCMC-OGFP ESC line at different time points of CHIR treatment. (G) (Top) Experimental scheme to analyse the ability of the Cdx2 or Oct4 or Cdx2+Oct4 expressing subpopulation of TCMC-OGFP cells to retain or lose the expression of Cdx2 or Oct4 in SL and TSC culture media. The TCMC-OGFP were cultured in SLCHIR for 48hrs and the cells were sorted as Cdx2 mCherry (P1), Cdx2-mCherry + Oct4-GFP (P2), and Oct4-GFP (P3) populations. The sorted cells were cultured in ESC culture media (SL) and TSC media (Tanaka et. al.2005). The cells were analysed for expression of CDX2 and OCT4 by immunostaining after 72 hours of culture. Scale bar = 100µm. (Bottom) Immunofluorescence assay for detection of CDX2 and OCT4 of P1, P2, and P3 subpopulation cells cultured in SL and TSC media. (H) Population of cells expressing CDX2 or OCT4 in subpopulations P1, P2, and P3 after 72hrs of culture in SL or TSC media. ‘*’ indicates a p-value <0.05, ‘**’ indicates a p-value <0.01, ‘***’ indicates a p-value <0.001, ‘****’ indicates a p-value <0.0001 ‘ns’ indicates a p-value > 0.05.

**Supplementary Fig. 2.**
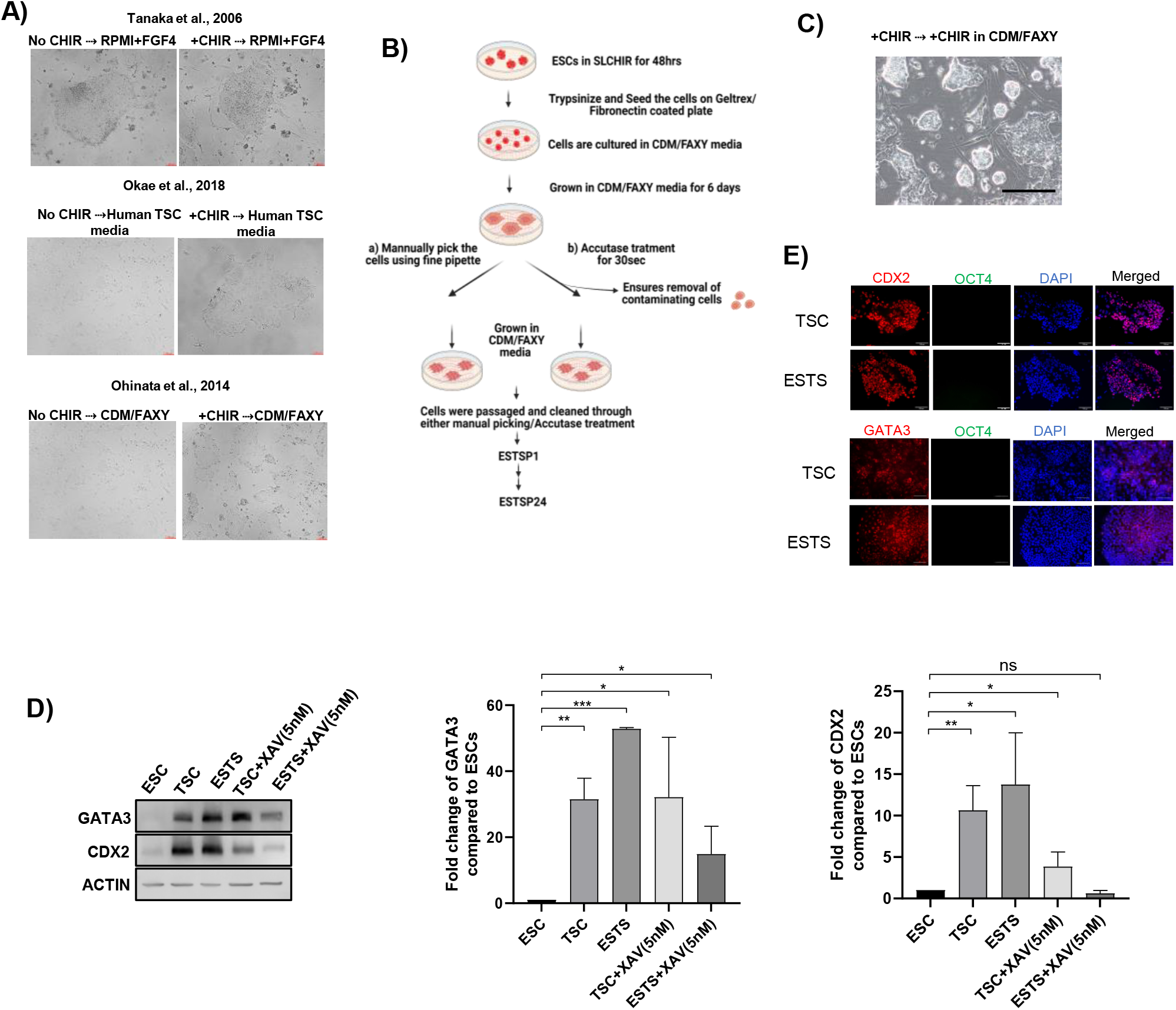
(A) Bright-field images of ESCs grown under indicated TSC culture conditions. Scale bar =100µm. (B) Flow chart showing the derivation of ESTS from ESC. (C) Phase contrast image of ESCs grown in continuous CHIR treatment in CDM/FAXY media. Scale bar = 100µm. D) (Left) immunoblot analysis of CDX2, GATA3, and ACTIN in TSC and ESTS cells in FAXY containing 5 nM XAV. (Right) Relative CDX2 and GATA3 levels as estimated by densitometry. The error bar represents the SD of the mean of biological replicates (n=3). ‘*’ indicates a p-value <0.05, ‘**’ indicates a p-value <0.01. (E) (Top) Immunofluorescence of CDX2 and OCT4 in TSC and ESTS cultured in CDM/FAXY. (Bottom) Immunofluorescence of GATA3 and OCT4 in TSC and ESTS cultured in CDM/FAXY media. Scale bar = 100µm.

**Supplementary Fig. 3.**
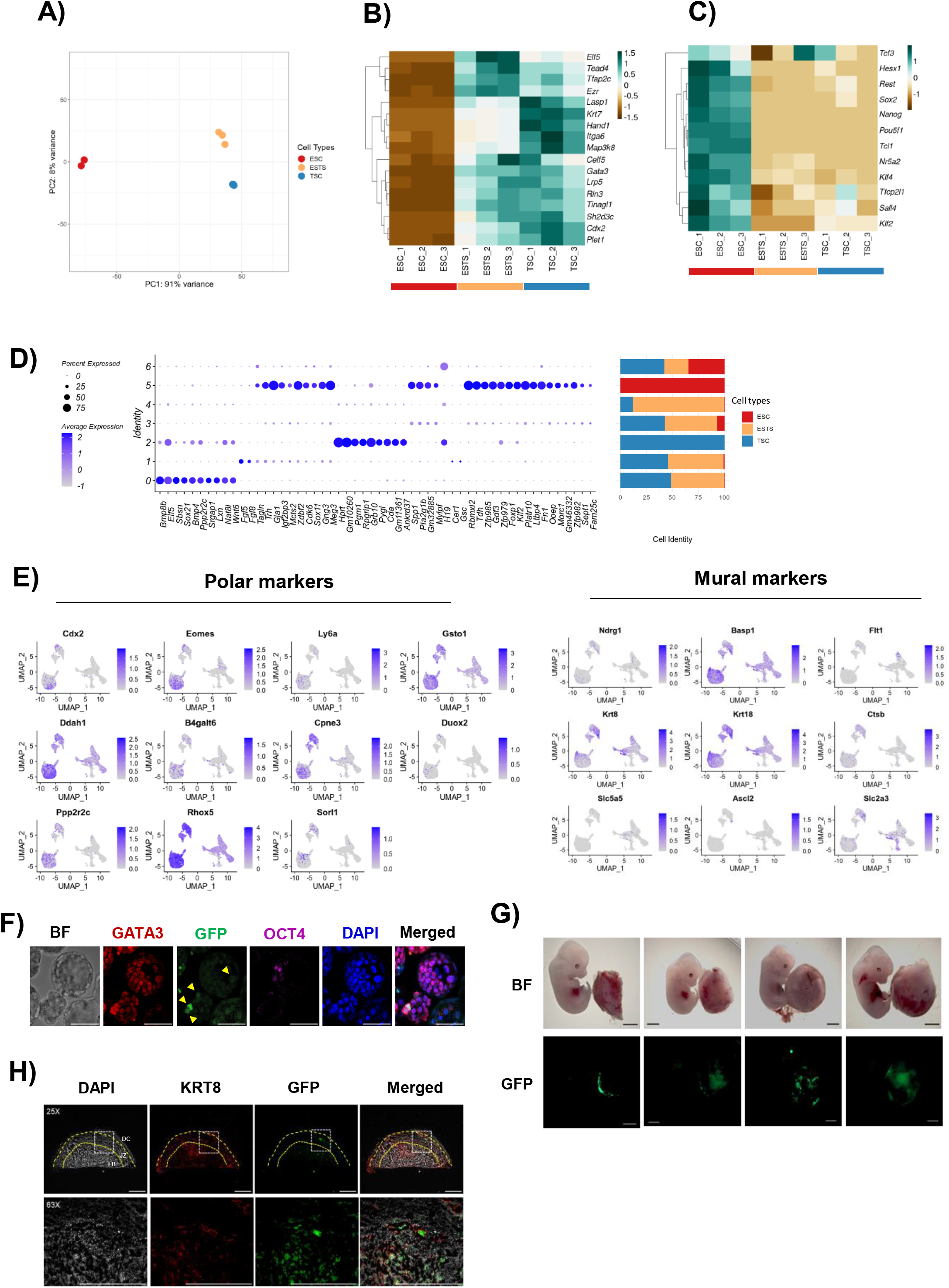
(A) Principal component analysis (PCA) of RNA-seq of ESC, TSC, and ESTS (B) Heatmap of expression of TE lineage genes from RNA-seq data of ESC, TSC, and ESTS. (C) Heatmap of expression of TE lineage genes from RNA-seq data of ESC, TSC, and ESTS (D) scater plot of the top 10 genes differentially expressed in different clusters of ESC, TSC, and ESTS from sc-RNA seq. (E) Feature plots projecting expression of representative polar and mural markers of TE overlaying in Fig 2A UMAP. (F) Immunofluorescence of GATA3, GFP and OCT4 in E3.5 after injection of GFP labelled ESTS (ESTS-GFP) into 8-cell stage embryo. Scale bar = 50µm (G) Representative image showing the contribution of GFP labelled ESTS descendant cells to the placenta at E12.5 (H) Immunofluorescence of KRT8 and GFP in the E12.5 placental section of ESTS transferred embryo. DC, JZ and LB depict the decidual region, junctional zone, and labyrinth region of the placenta respectively. Scale bar = 2000µm

**Supplementary Fig. 4.**
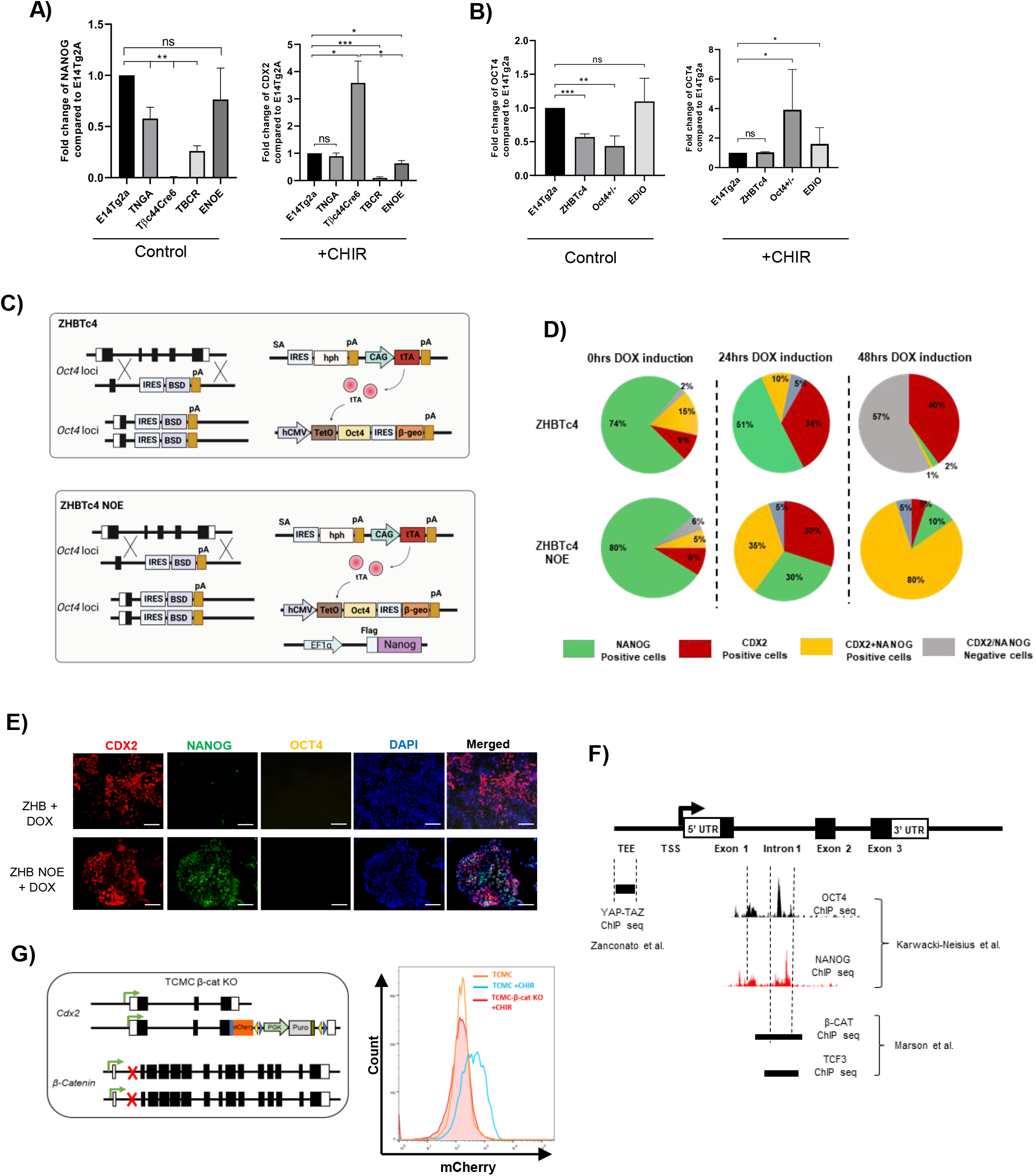
(A) Relative CDX2 and NANOG levels in cells cultured in SL and SLCHIR as estimated by densitometry of the indicated cell lines (n=3). B) Relative CDX2 and OCT4 levels in cells cultured in SL and SLCHIR as estimated by densitometry of the indicated cell lines (n=3). ‘*’ indicates a p-value <0.05, ‘**’ indicates a p-value <0.01, ‘***’ indicates a p-value <0.001, ‘ns’ indicates a p-value > 0.05. (C) Schematic depiction of ZHBTc4 and ZHBTc4 NOE cell lines. (D) Pie chart representing the percentage of the cells expressing CDX2 and NANOG upon doxycycline treatment in ZHB and ZHB NOE cell lines at a different time point of doxycycline treatment. (E) Immunofluorescence of CDX2, NANOG, and OCT4 in ZHB and ZHB NOE cell lines treated with or no Doxycycline. (Scale bar = 100µm). (F) Schematic depicting the analysis of ChIP enrichment peaks previously pulled down by OCT4, NANOG, YAP/TAZ, β-CATENIN, and TCF3 on Cdx2 loci by different experiments and groups listed. (G) (Left) Schematic depiction of TCMCβ-Catenin-/-. The cell line was derived from TCMC by CRISPR-mediated knockout of β-Catenin. (Right) histogram profile of mCherry expression in TCMC, TCMC treated with CHIR, and TCMCβ-Cat -/- treated with CHIR.

**Supplementary Fig. 5.**
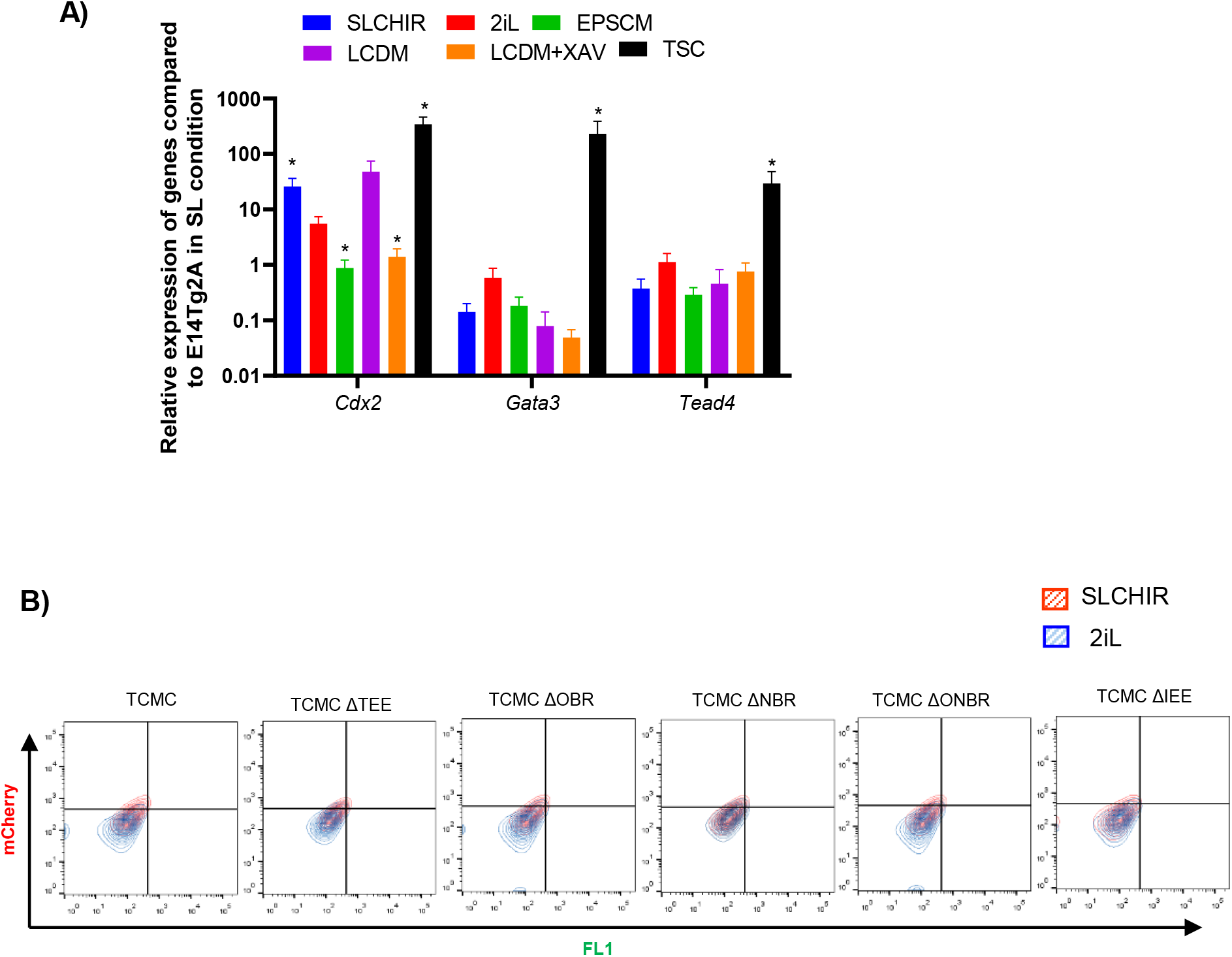
(A) Relative expression levels of indicated TE lineage genes Cdx2, Gata3, and Tead4 in ESC cultured in the indicated culture conditions. TSCs were used as a positive control. (B) Contour plots generated from FACS and showing the expression of mCherry in TCMC, TCMCΔTEE, TCMCΔOBR, TCMCNBR, TCMCΔONBR, and TCMCΔIEE cultured in SL and 2iL.

## Key resource table

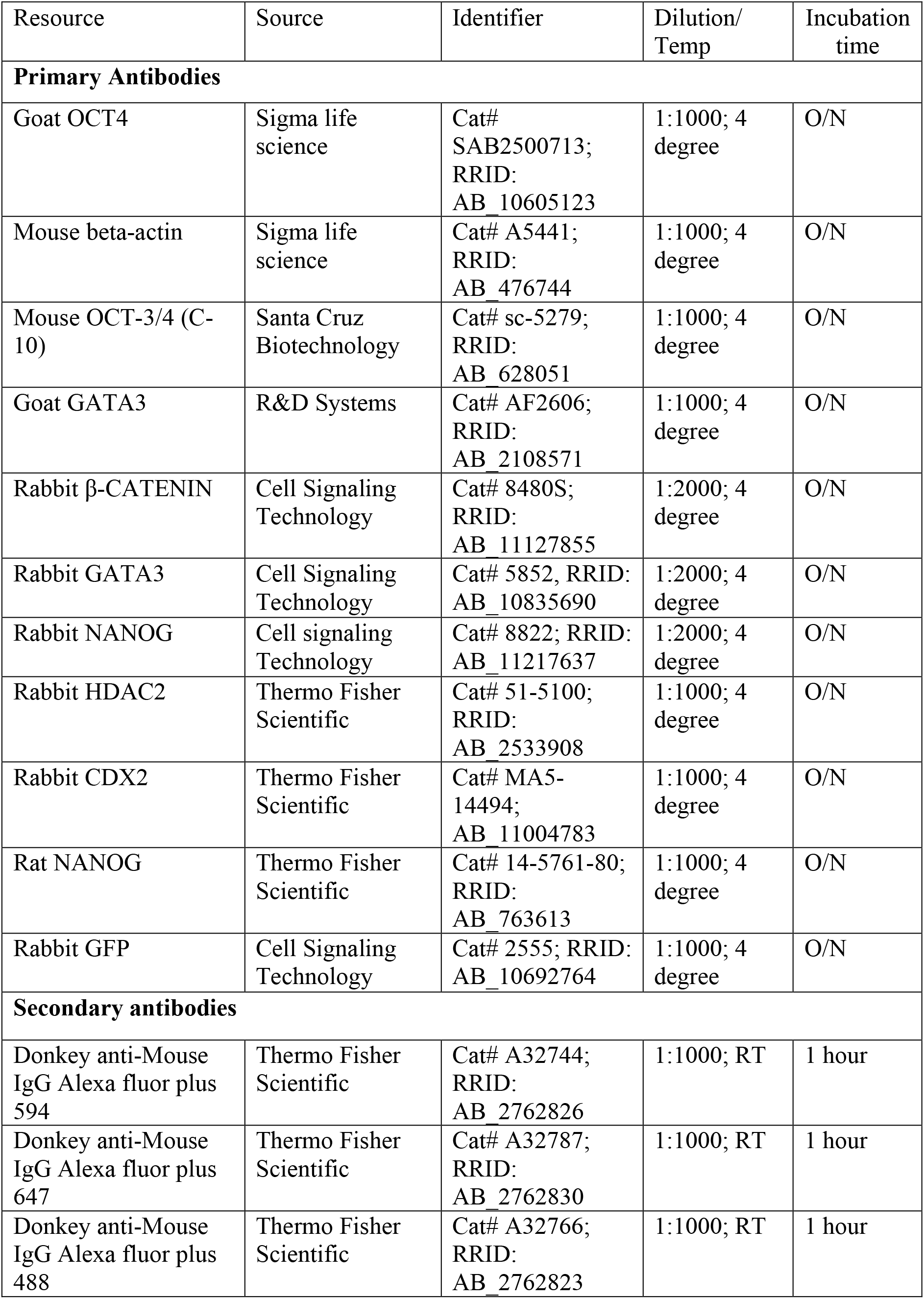

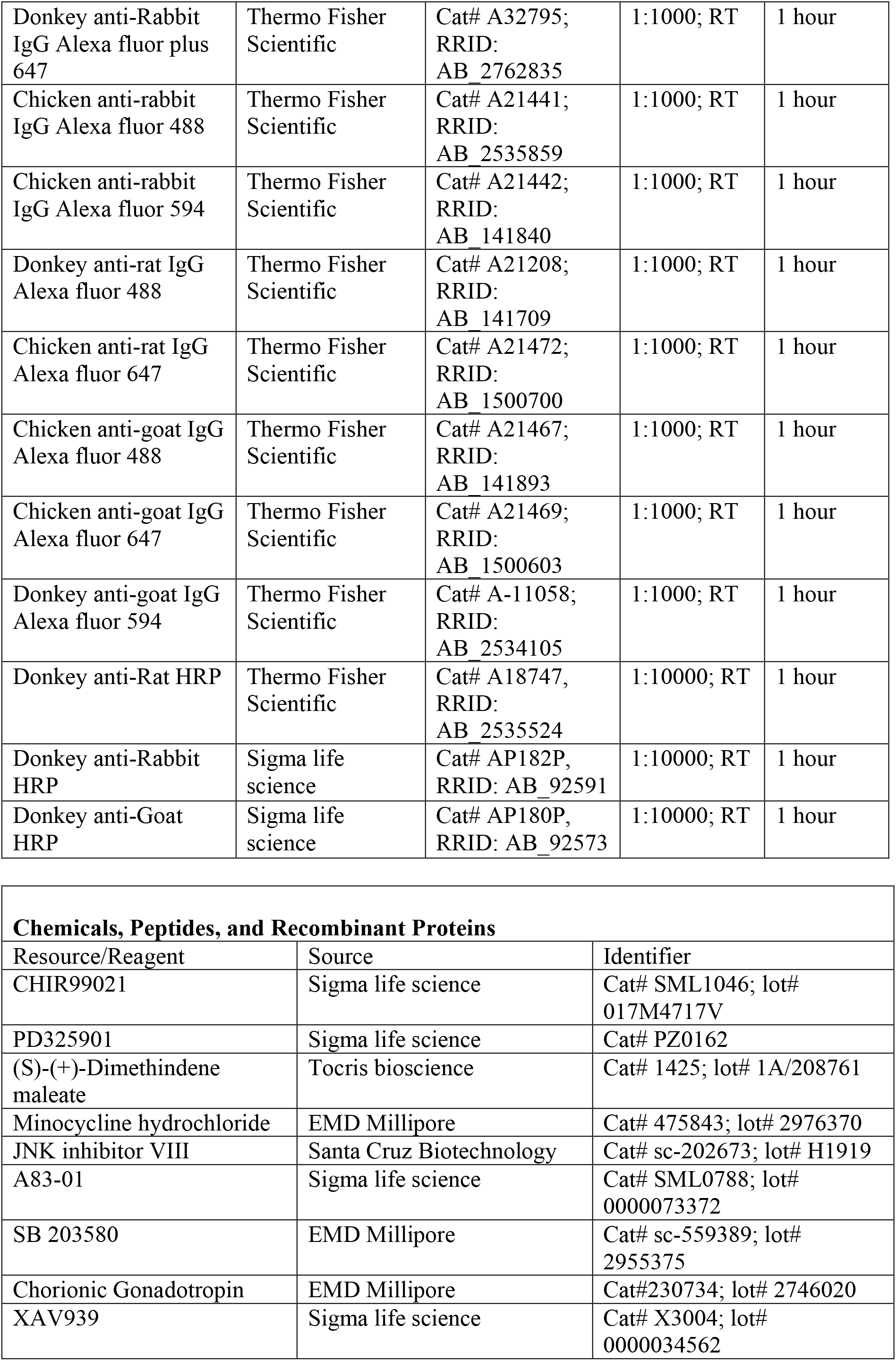

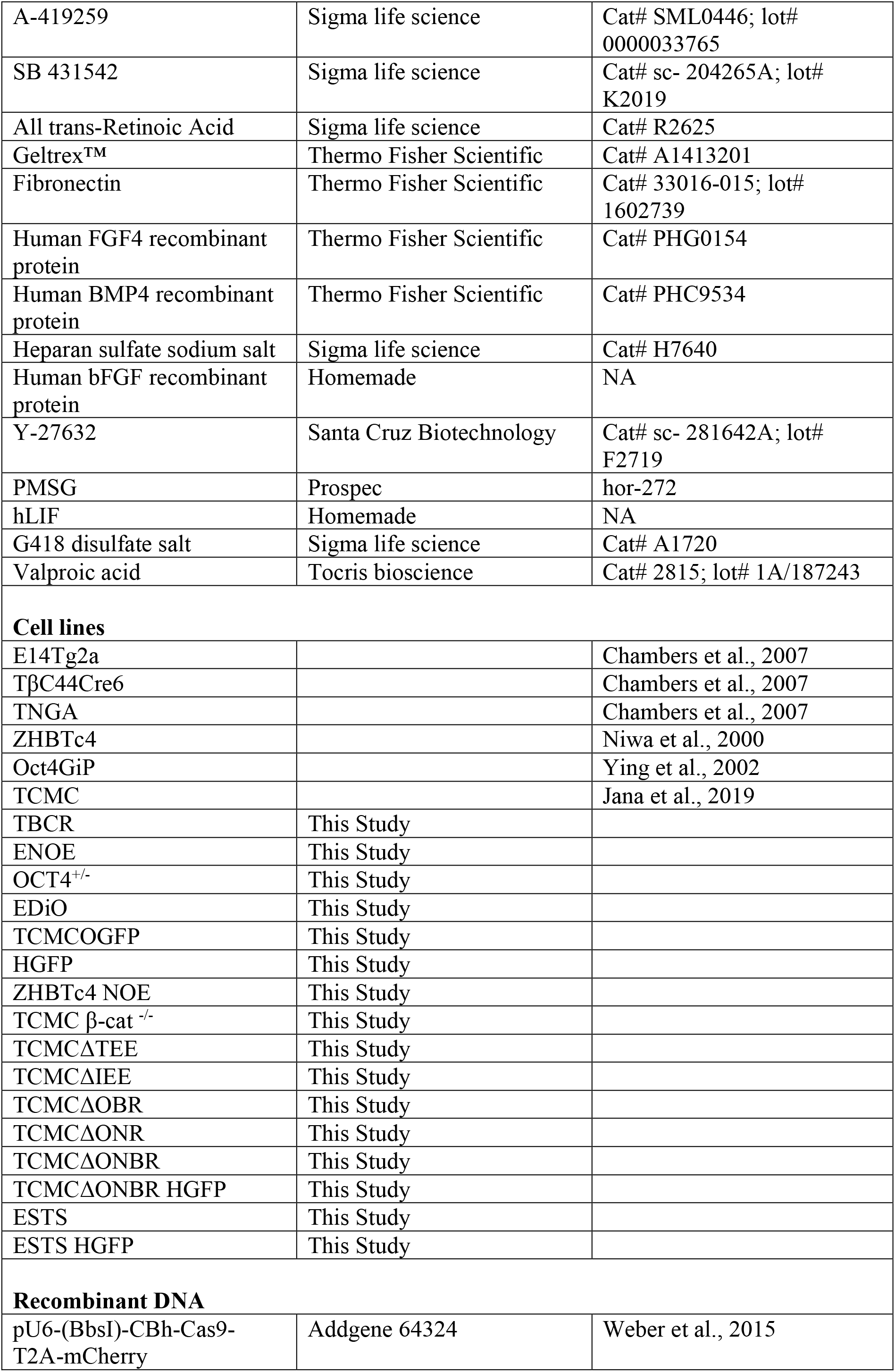

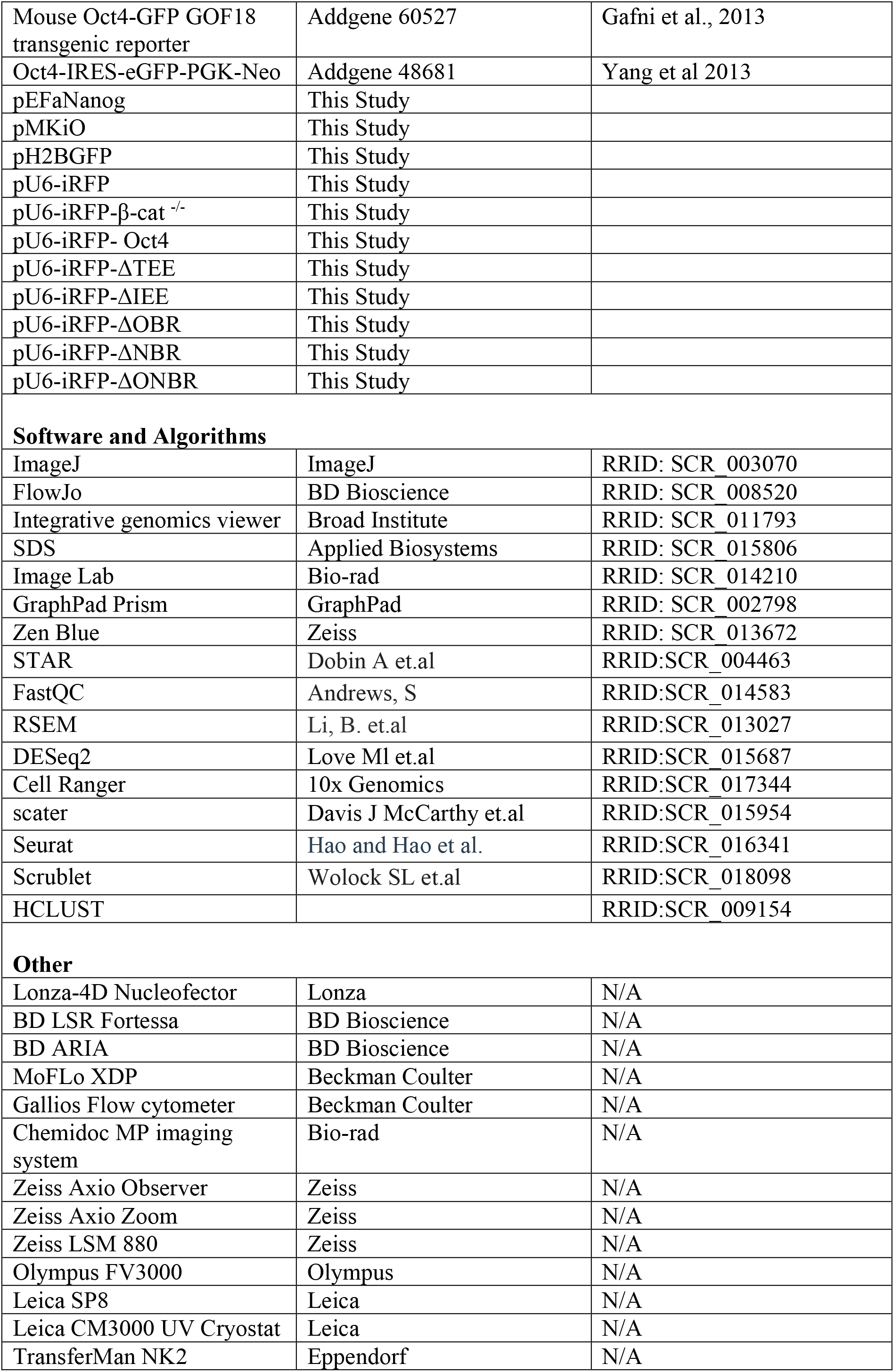

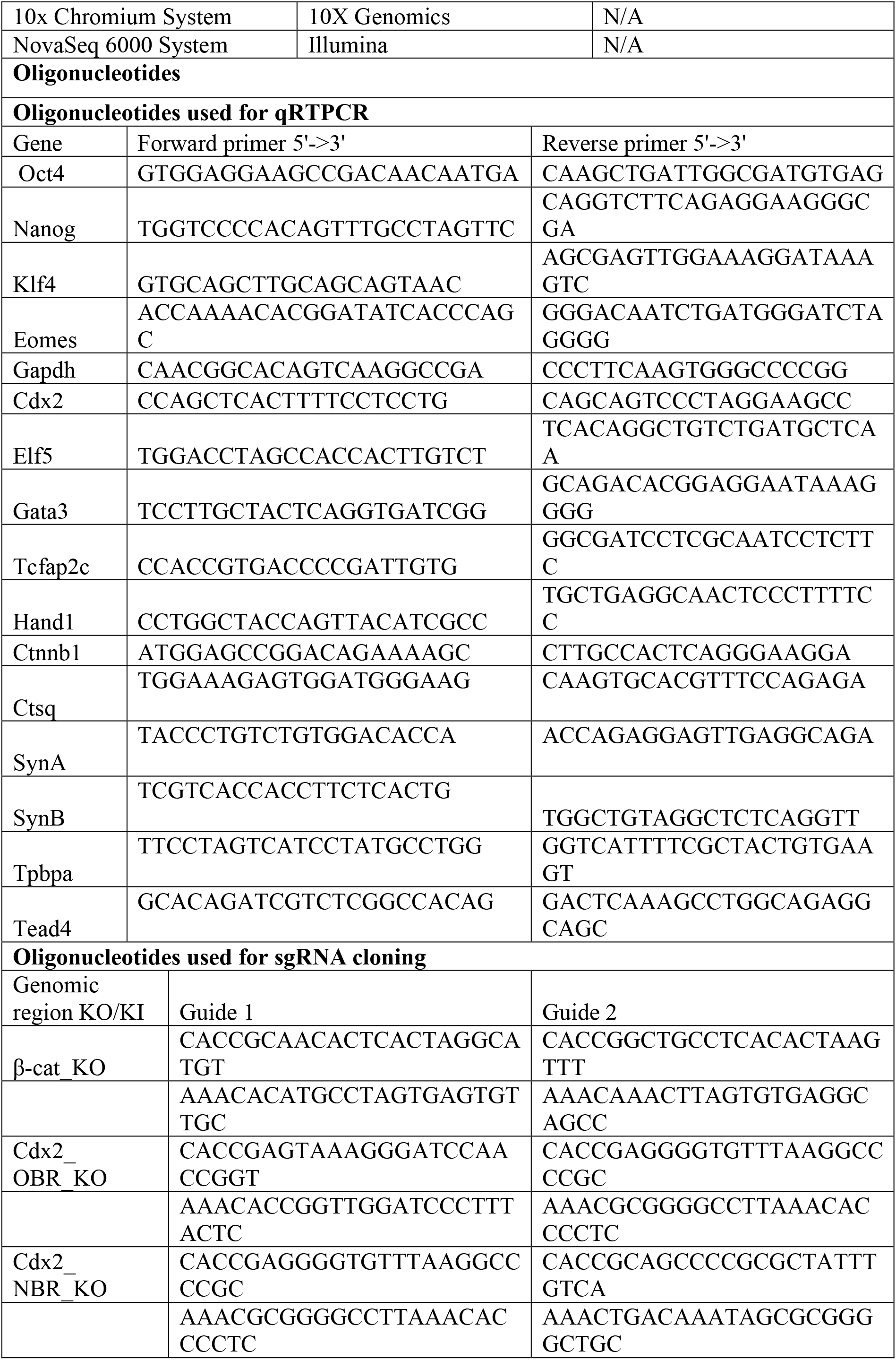

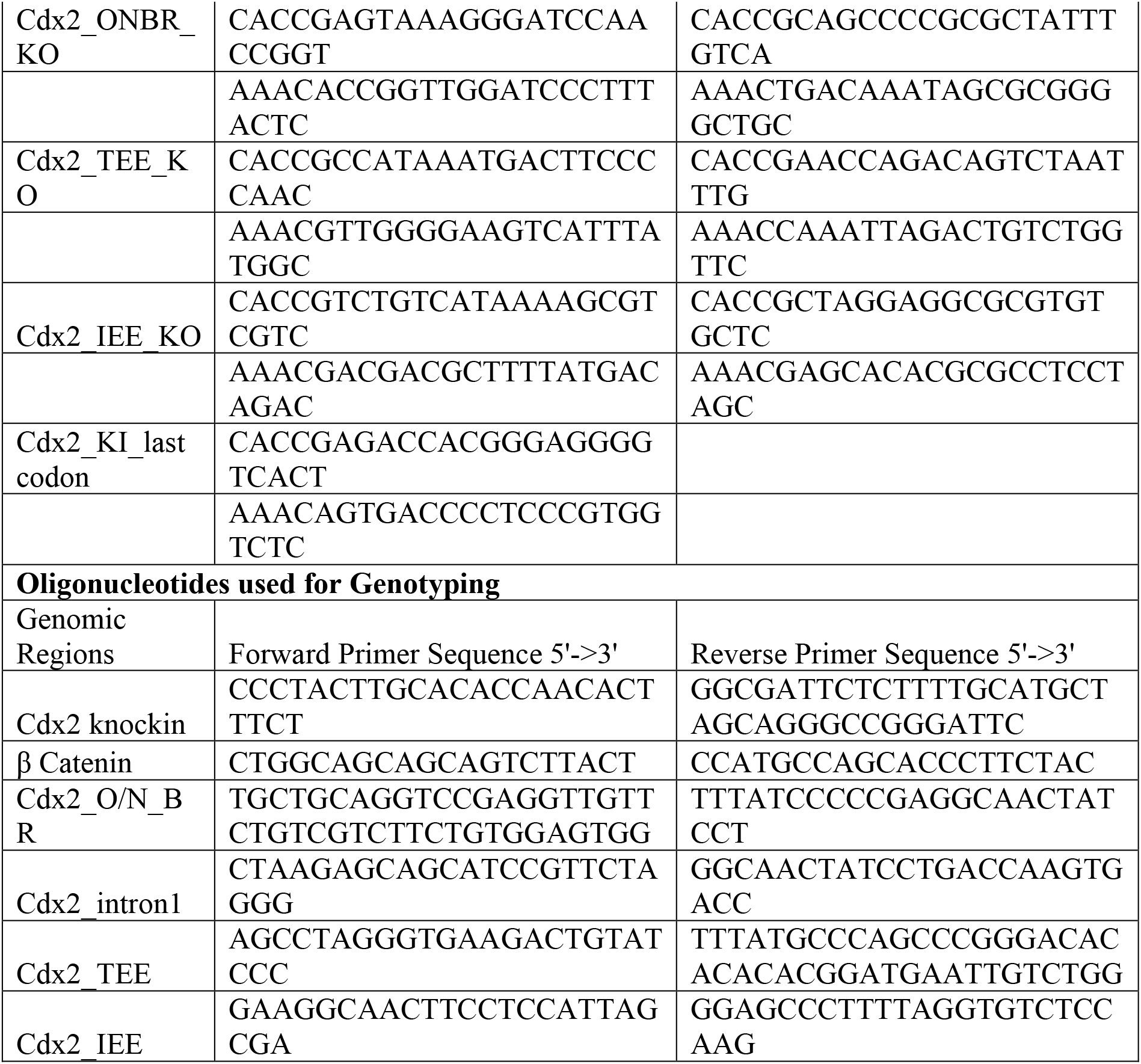

## Cell Pedigree Chart

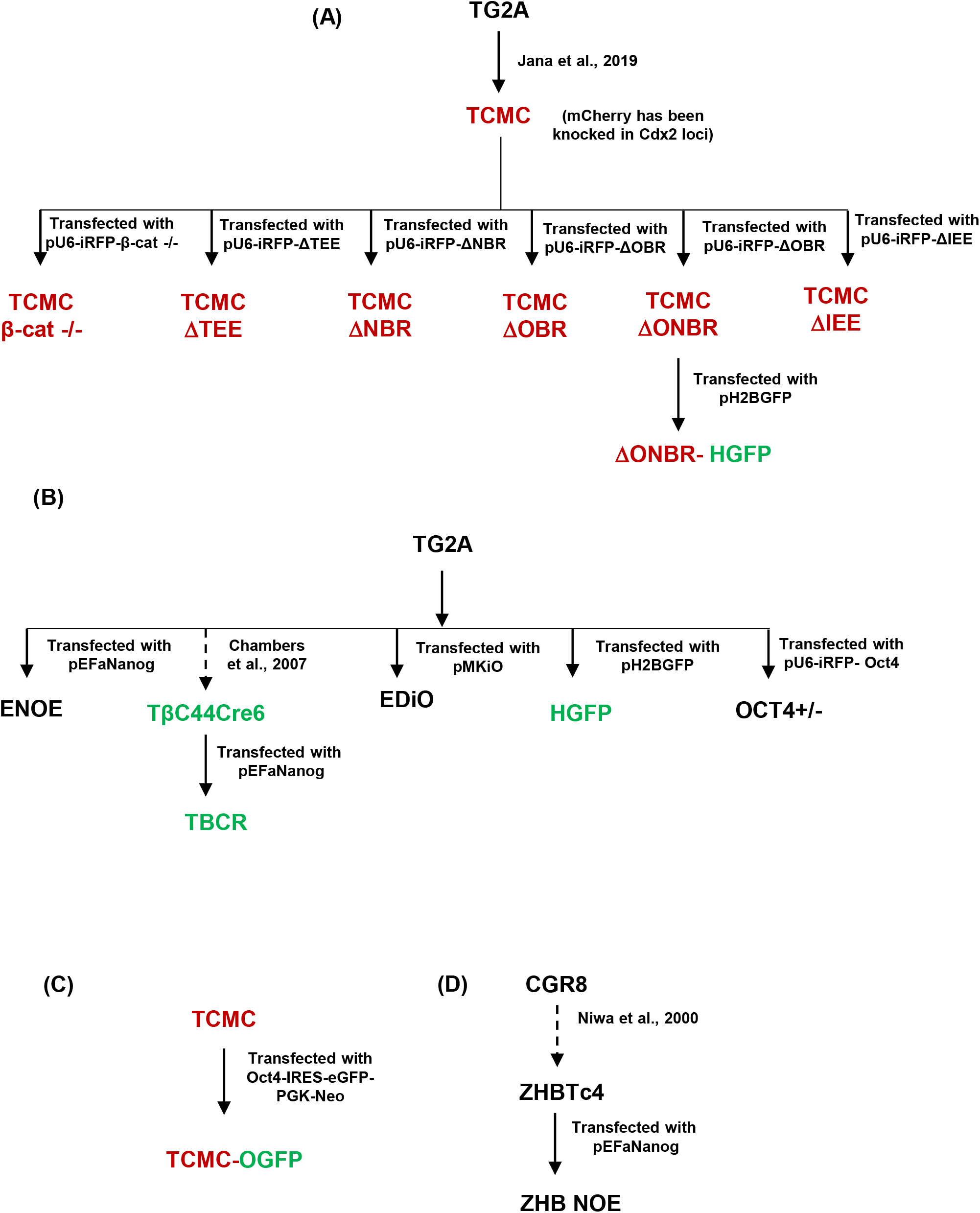

Figure: Information related to methods. A pedigree chart of cell lines used in this study low chart illustrating the lineage and process of generation of TCMC and generation of k out cells lines in TCMC background. (B) A flow chart Illustrating generation of neered lines in E14Tg2a background. (C) A flow chart depicting derivation of knock-in line TCMC-OGFP in TCMC background. (D) A flow chart depicting generation of ZHB E from ZHBTc4.

## Appendix 1

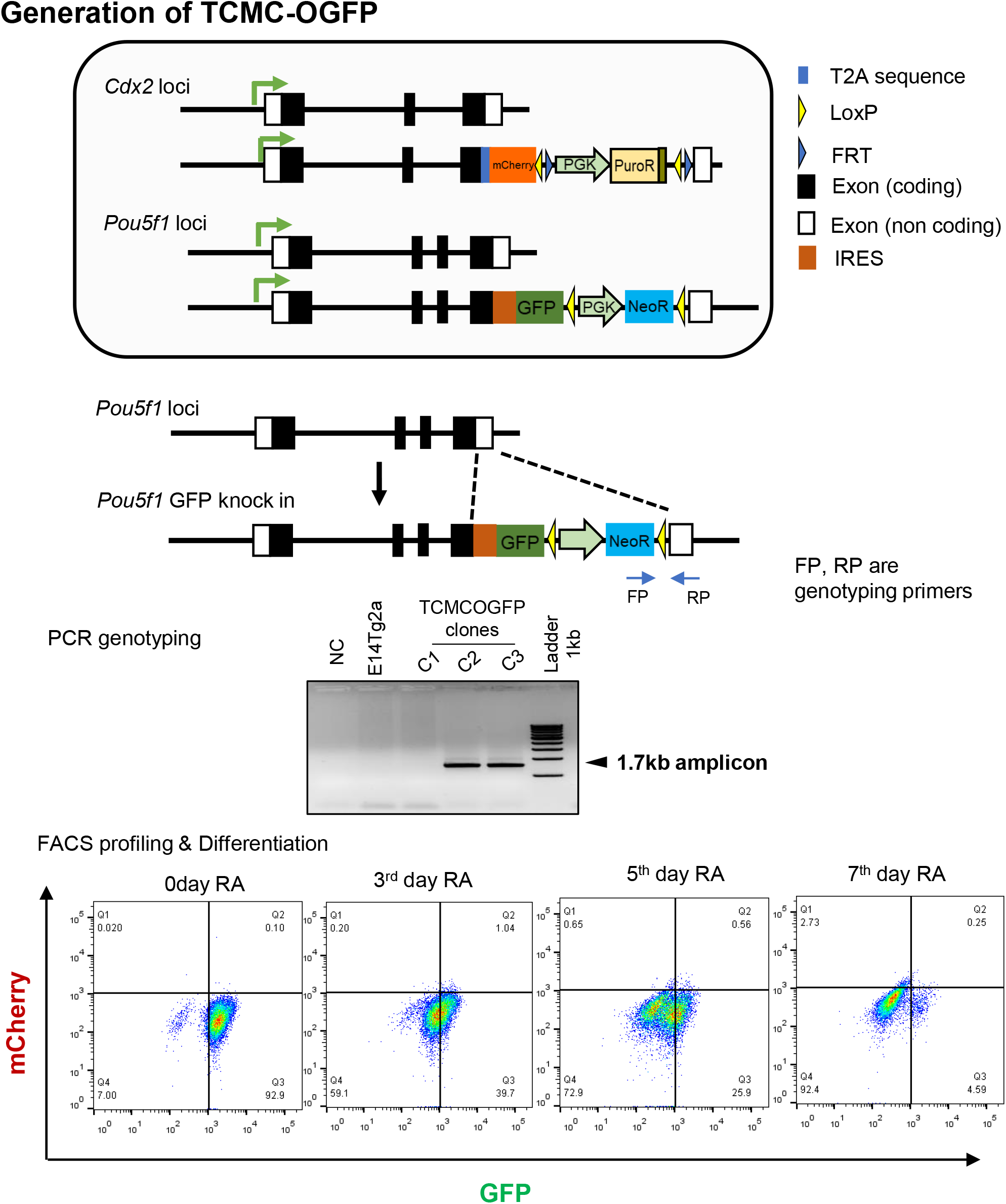

**Generation of TCMC-OGFP** (top) Schematic depiction of TCMC-OGFP cell line where one of e allele of *Oct4* has been knocked in with GFP and one of the allele of *Cdx2* has been ocked in with mCherry. The cell line has been derived from TCMC (middle) GFP knock in ct4 loci has been confirmed by genotyping and sequencing. (bottom) FACS profile of GFP pression was used as a functional reporter for the knock-in GFP in *Oct4* loci with TCMC-GFP cells grown in differentiating media with Retinoic acid (RA)

## Appendix 2

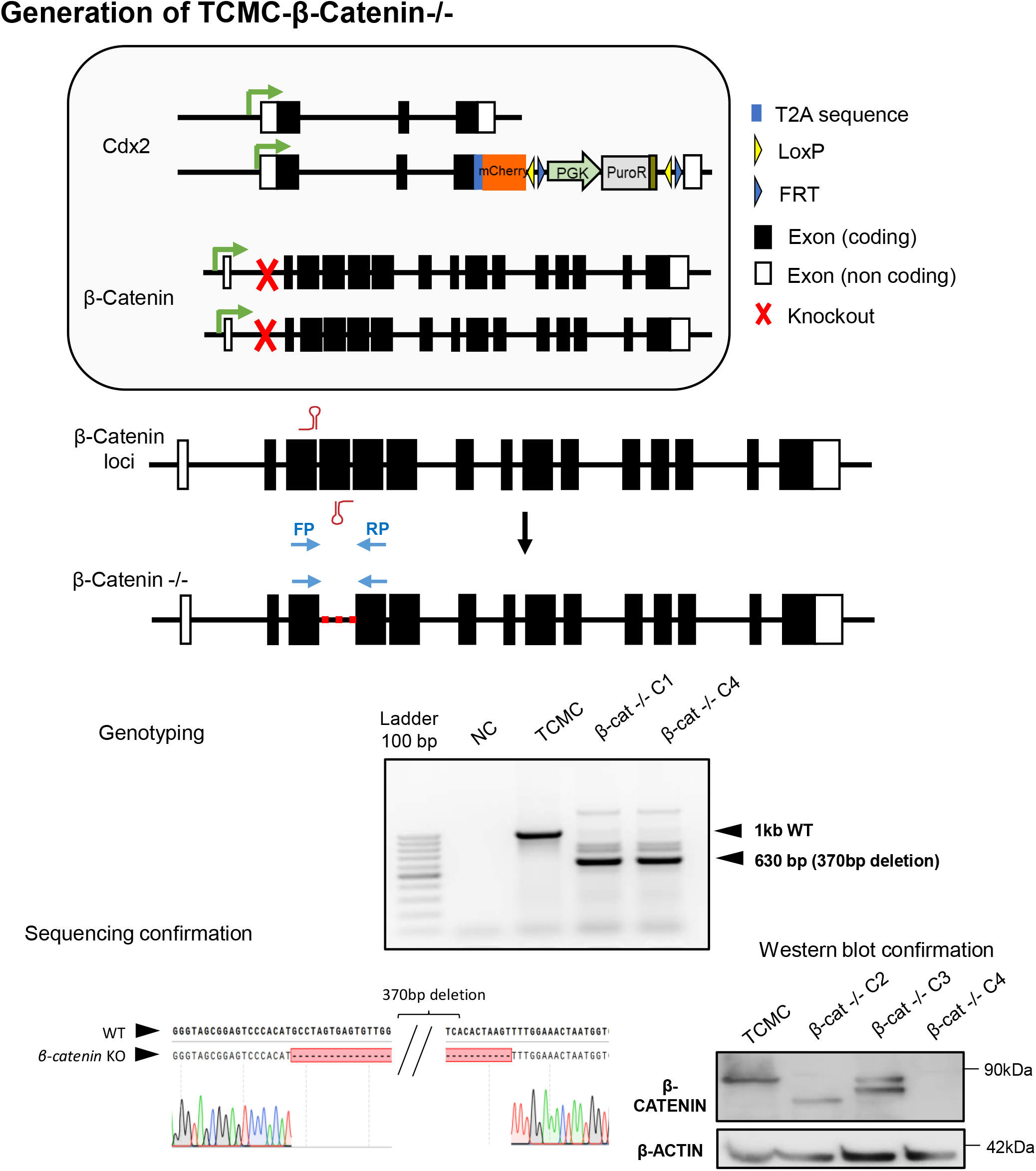

**Generation of TCMC-β-Catenin-/- ES cell line**(top) Schematic depiction of TCMC-β-catenin ^-/-^ cell line. The cell line has been derived from TCMC (middle) β-catenin ^-/-^ has been confirmed enotyping encompassing the deletion site. and sequencing. (bottom) Western blotting for rent clones were confirmed by checking β-CATENIN.

## Appendix 3

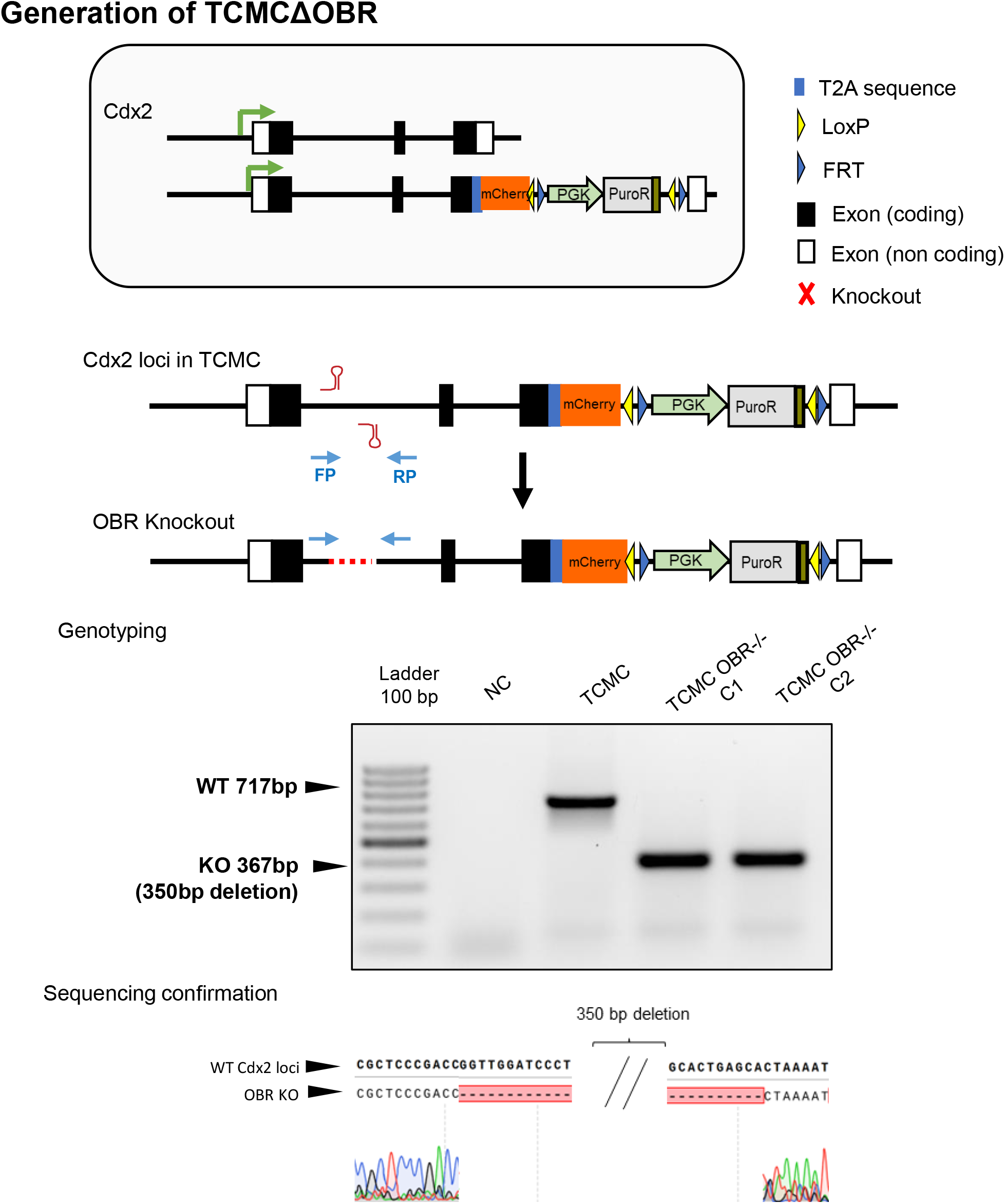

**Generation of TCMCΔOBR** (top) Schematic depiction of TCMCΔOBR The cell line has been derived from TCMC (middle) ΔOBR has been confirmed by genotyping different clones encompassing the deletion site. (bottom) Sequencing confirmation of the genotyped PCR product.

## Appendix 4

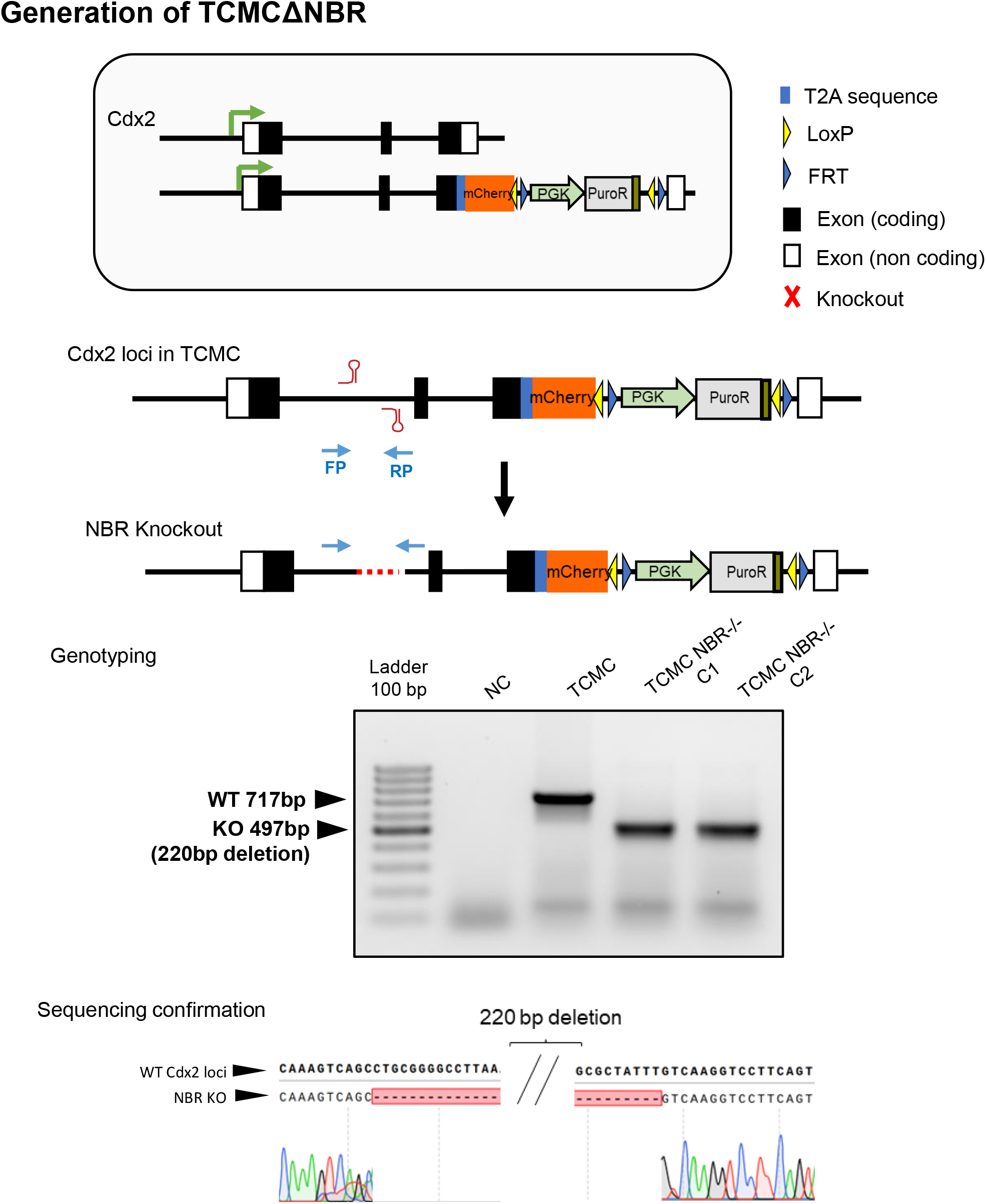

**Generation of TCMCΔNBR** (top) Schematic depiction of TCMCΔNBR The cell line has been derived from TCMC (middle) ΔNBR has been confirmed by genotyping different clones encompassing the deletion site. (bottom) Sequencing confirmation of the genotyped PCR product.

## Appendix 5

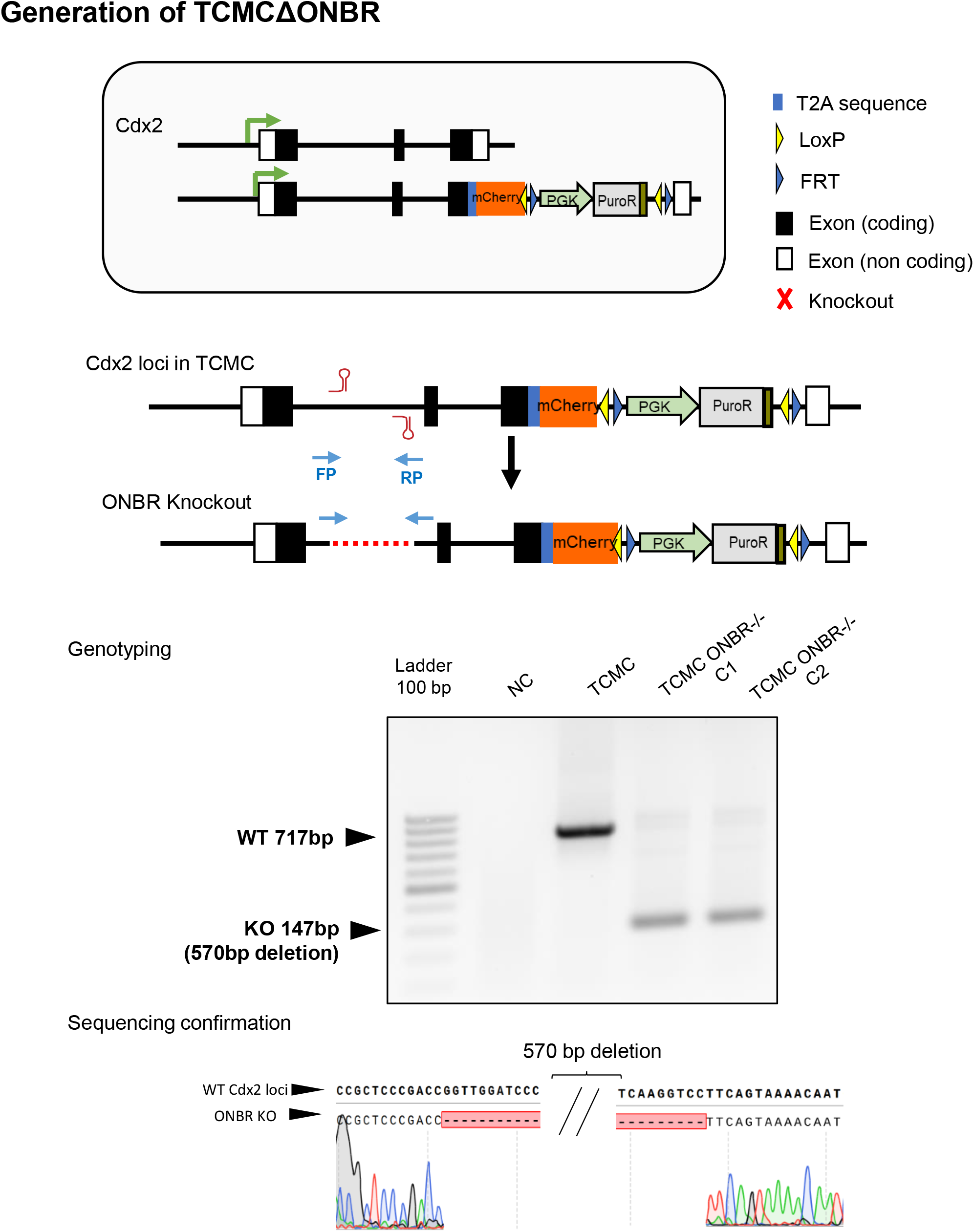

**Generation of TCMCΔONBE** (top) Schematic depiction of TCMCΔONBR The cell line has been derived from TCMC (middle) ΔONBR has been confirmed by genotyping different clones encompassing the deletion site. (bottom) Sequencing confirmation of the genotyped PCR product.

## Appendix 6

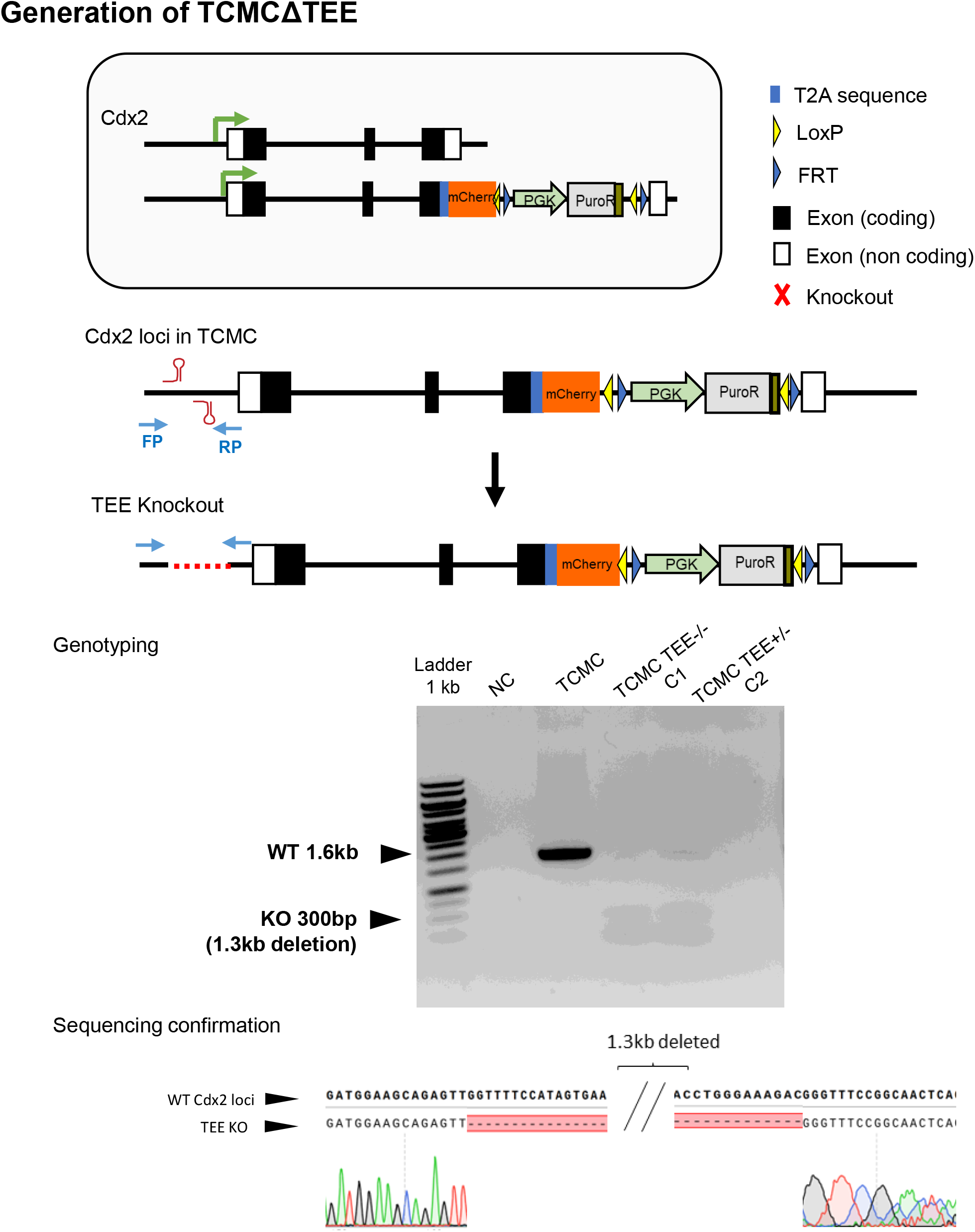

**Generation of TCMCΔIEE** (top) Schematic depiction of TCMCΔTEE The cell line has been derived from TCMC (middle) ΔTEE has been confirmed by genotyping different clones encompassing the deletion site. (bottom) Sequencing confirmation of the genotyped PCR product.

## Appendix 7

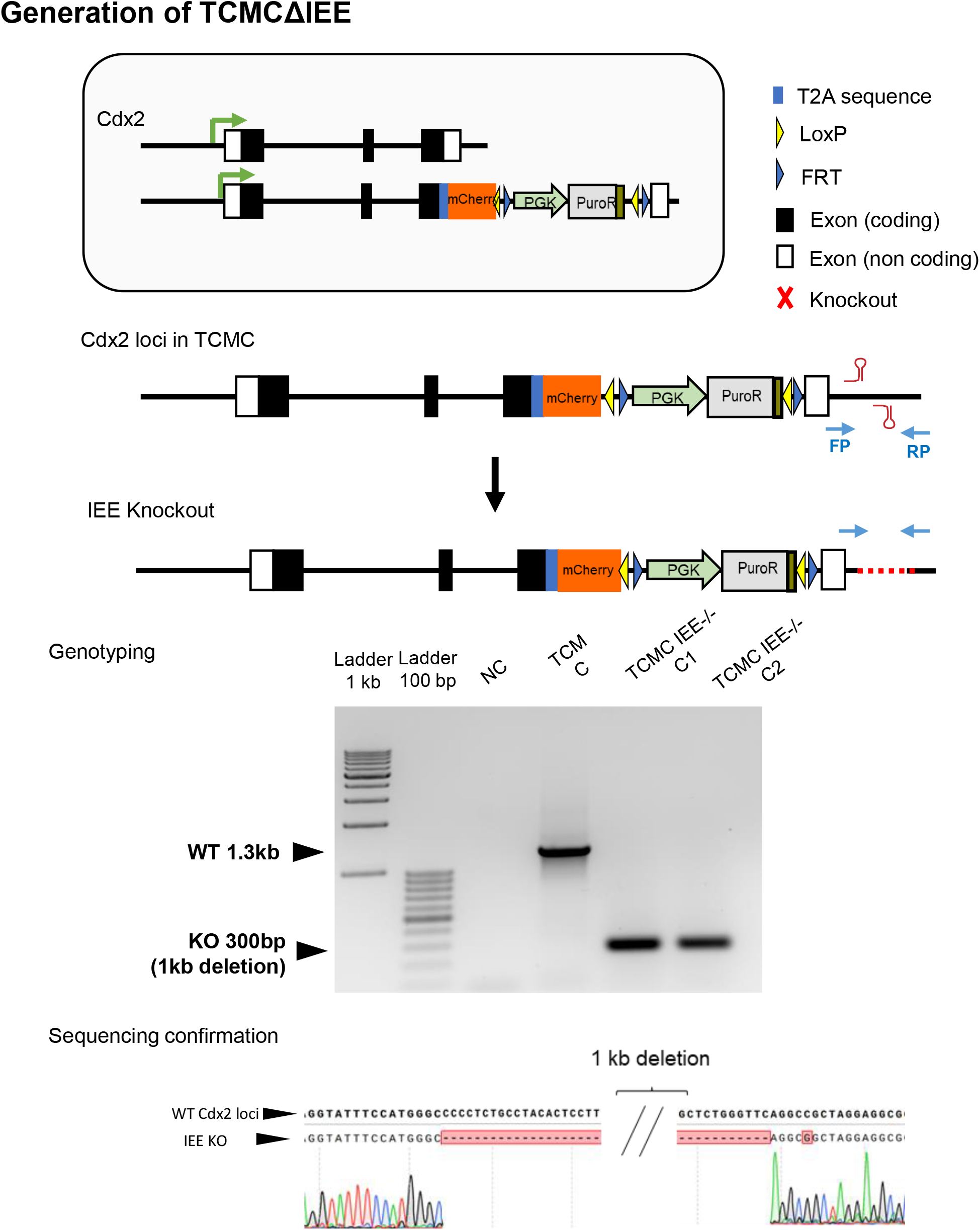

**Generation of TCMCΔIEE** (top) Schematic depiction of TCMCΔIEE The cell line has been derived from TCMC (middle) ΔIEE has been confirmed by genotyping different clones encompassing the deletion site. (bottom) Sequencing confirmation of the genotyped PCR product.

## Notes

### Competing Interest Statement

The authors have declared no competing interest.

### Summary of Updates

Error in line no. 499 in the method section, the final concentration of XAV and Y27632 has been corrected.

